# Compressive Pangenomics Using Mutation-Annotated Networks

**DOI:** 10.1101/2024.07.02.601807

**Authors:** Sumit Walia, Harsh Motwani, Kyle Smith, Russell Corbett-Detig, Yatish Turakhia

## Abstract

Pangenomics is an emerging field that uses a collection of genomes of a species instead of a single reference genome to overcome reference bias and study the within-species genetic diversity. Future pangenomics applications will require analyzing large and ever-growing collections of genomes. Therefore, the choice of data representation is a key determinant of the scope, as well as the computational and memory performance of pangenomic analyses. Current pangenome formats, while capable of storing genetic variations across multiple genomes, fail to capture the shared evolutionary and mutational histories among them, thereby limiting their applications. They are also inefficient for storage, and therefore face significant scaling challenges. In this manuscript, we propose PanMAN, a novel data structure that is information-wise richer than all existing pangenome formats – in addition to representing the alignment and genetic variation in a collection of genomes, PanMAN represents the shared mutational and evolutionary histories inferred between those genomes. By using “evolutionary compression”, PanMAN achieves 5.2 to 680-fold compression over other variation-preserving pangenomic formats. PanMAN’s relative performance generally improves with larger datasets and it is compatible with any method for inferring phylogenies and ancestral nucleotide states. Using SARS-CoV-2 as a case study, we show that PanMAN offers a detailed and accurate portrayal of the pathogen’s evolutionary and mutational history, facilitating the discovery of new biological insights. We also present *panmanUtils*, a software toolkit that supports common pangenomic analyses and makes PanMANs interoperable with existing tools and formats. PanMANs are poised to enhance the scale, speed, resolution, and overall scope of pangenomic analyses and data sharing.

## Introduction

Pangenomics, the area of bioinformatics that deals with representing a collection of genomes from a single species, has gained significant interest in recent years^1–3^. Interest is driven by the dramatic advancements in genome sequencing technologies over the past two decades which have drastically reduced costs and increased throughput, enabling the research community to sequence and examine a vast number of genomes within the same species. For example, during the recent COVID-19 pandemic, over 16 million genomes of SARS-CoV-2 alone were sequenced globally and shared on online databases^4,5^. This wealth of data enabled the rapid identification of new SARS-CoV-2 variants^6,7^, monitoring the prevalence of circulating variants^8^, assessing the fitness of different variants^9^, investigating the transmission patterns of outbreaks^10,11^, and aiding in the development of vaccines optimized for specific variants^12,13^. Because they represent the full range of known genetic variation, pangenomes also mitigate the problems of “reference bias” that have impacted biomedical research due to the traditional dependence on a single linear reference sequence for each species^14–19^.

The data structures used for pangenomics research are critical because they not only determine the efficiency of the representation but also the representative power itself. Graph-based pangenome formats^18,20–27^, also sometimes referred to as “graph genomes”, have become popular and widely adopted, largely because of the interest in reducing the reference bias in read mapping and variant calling applications. However, graph genomes only represent the genetic variation in a collection of genomes, but not their shared evolutionary and mutational histories, which limits the scope of their application. They also have large storage needs which typically do not scale well with the number of sequences. Recent proposals, such as AGC^28^ and MiniPhy^29^, improve the storage efficiency of pangenomes by extending traditional data compression algorithms to genome sequences. While these methods provide excellent compression, their utility is limited to storage and fast retrieval of sequences, as they do not explicitly store variation or any evolutionary information. GBZ^21^ is another format proposed recently to compress graph genomes. However, despite offering substantial improvements in storage efficiency, GBZ also suffers from the same limitations in terms of representative power.

Because of the aforementioned limitations of graph genomes, new formats represent evolutionarily relevant information, such as the phylogenetic relationships among samples and inferred occurrences of mutations, in large collections of genomes for specific applications. Two examples include the mutation-annotated tree (MAT) in UShER^30^ and tree sequences in tskit^31^. UShER-MAT stores a phylogenetic tree corresponding to the sequences in the collection with sequences at the tips and branches annotated with substitutions that are inferred to have occurred on them. UShER-MAT introduced the notion of “evolutionary compression” and was widely adopted during the COVID-19 pandemic for analysis and data sharing, establishing itself as the *de facto* standard tool for naming SARS-CoV-2 variants^7^. However, UShER-MAT is a lossy format, i.e., it cannot reconstruct the original sequences from which it was derived, since it captures only single nucleotide substitutions, and excludes insertions, deletions, and rearrangements. Likewise, tree sequences in tskit store a series of trees to depict the distinct evolutionary pathways of the different genomic regions that have undergone recombination, a simplified ancestral recombination graph (ARG)^32–36^. The format also has the ability to store all the information in a Variant Call Format (VCF) file. Still, like VCF, it lacks the ability to record nested mutations, and is, therefore, lossy. Fig. 1 summarizes the representative power and limitations of the existing pangenomic data structures.

**Figure 1:**
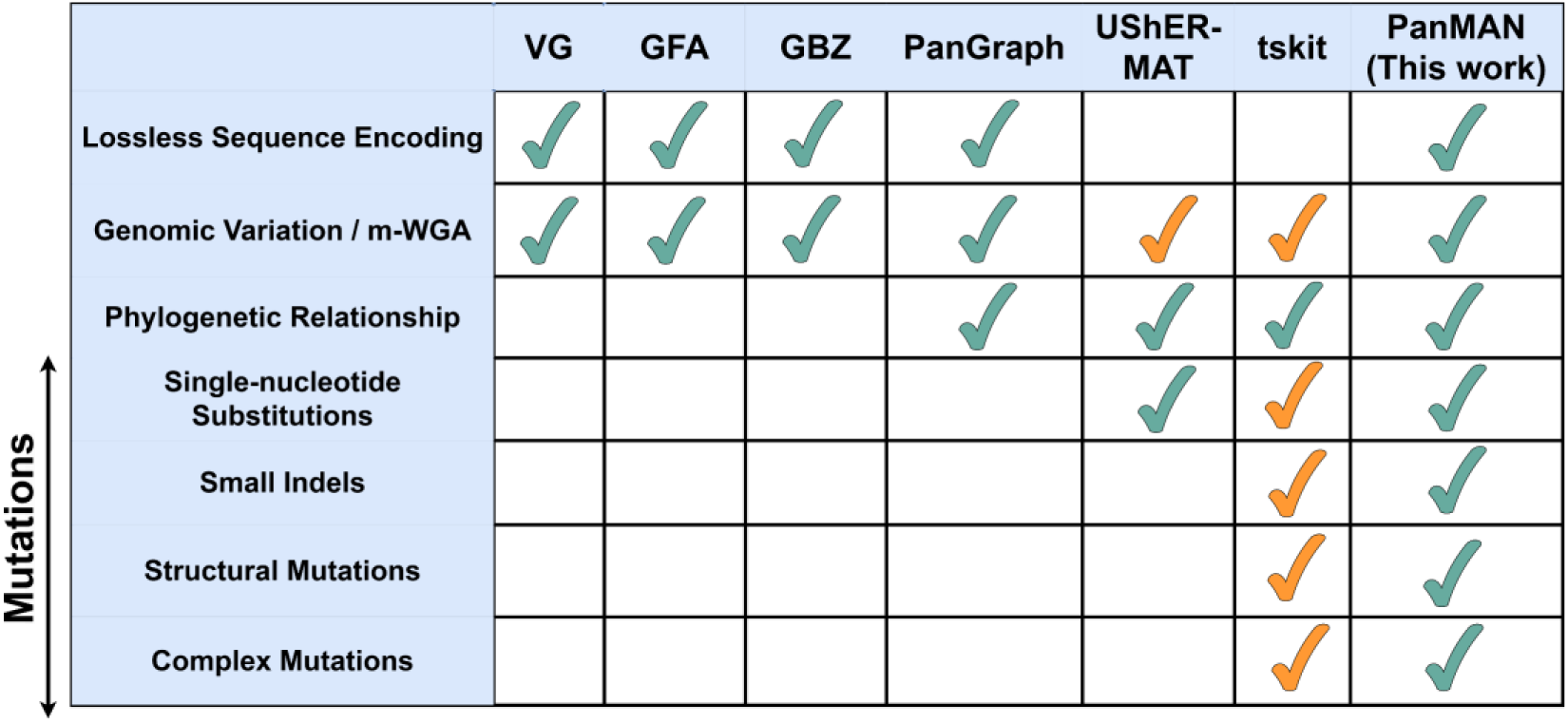
Comparison of representative power of PanMAN against other pangenomic formats. (yellow ticks indicate partial representative ability).

In this manuscript, we propose Pangenome Mutation-Annotated Networks (PanMANs), a novel data representation for pangenomes that provides massive leaps in both representative power and storage efficiency. PanMANs use evolutionary compression as in UShER-MAT. Specifically, PanMANs are composed of mutation-annotated trees, called PanMATs, which, in addition to substitutions, also annotate inferred insertions and deletions on the different branches. Multiple PanMATs are connected in the form of a network using edges to generate a PanMAN. These edges store recombination and horizontal gene transfer, which result in sequences involving multiple parent sequences and violate the vertical inheritance assumption of single trees. We found PanMANs to be the most compressible of existing variation-preserving pangenome formats for the different microbial datasets we evaluated. We demonstrate PanMAN representative capabilities using the example of SARS-CoV-2 and show that PanMANs provide a more detailed and accurate representation of the pathogen’s evolutionary and mutational histories. Notably, PanMAN represents indels and recombination events that are often ignored in bioinformatic analyses^33,37^. We also demonstrate that PanMANs are compatible with any phylogenetic and ancestral state reconstruction algorithms. To make PanMANs more accessible and interoperable, we also developed a software utility, called *panmanUtils*, that supports functionalities to construct, modify, and extract useful information from PanMANs, including mutations, phylogeny, multiple whole-genome alignments (m-WGA), and genetic variations.

Overall, PanMANs have the potential to improve the scale, speed, and resolution of microbial pangenomic analysis, leading to new applications and insights.

## Results

### An Overview of the PanMAN Data Structure

PanMAN is designed to provide a lossless encoding of a collection of sequences and represent the complex phylogenetic and mutational histories shared between them (Fig. 1). In a subsequent subsection, we will discuss how PanMAN implicitly preserves the multiple whole-genome alignment (m-WGA) along with the genomic variation in the sequence collection, which can be exported using *panmanUtils*. At a high level, the PanMAN’s data structure comprises one or multiple *PanMATs*, each designed to encode sequences that are derived from a single ancestral sequence via vertical descent. PanMATs use a phylogenetic tree structure with a root sequence and branches annotated with mutations, allowing sequences at the tree’s tips to be derived by following these mutations from the root sequence, a storage-efficient method that was introduced in UShER^30^ as “evolutionary compression”. To manage indels and structural rearrangements, PanMAT introduces a three-level, reference-free coordinate system inspired by PanGraph^22^ (Fig. 2a-b, Methods). At the top level, this system uses *blocks* representing homologous or unique segments, which are each assigned a unique identifier based on their position in the linear ordering of blocks at the *pseudo-root* (Fig. 2a, Methods). The two lower levels in this coordinate system track the specific coordinates of nucleotides within blocks while handling small indels (Fig. 2b, Methods). Additionally, PanMAN can represent complex genetic events like recombination and horizontal gene transfer through network edges that connect nodes within and across PanMATs. The network edges record the complex mutations by storing the breakpoint coordinates in the parent sequences that underwent complex mutation and gave rise to the root of another PanMAT(Fig. 2c, Methods). The PanMAN data structure can also be stored as a file using serialization libraries (Methods). A more detailed explanation of PanMAN’s data structure and file format is provided in Methods.

**Figure 2:**
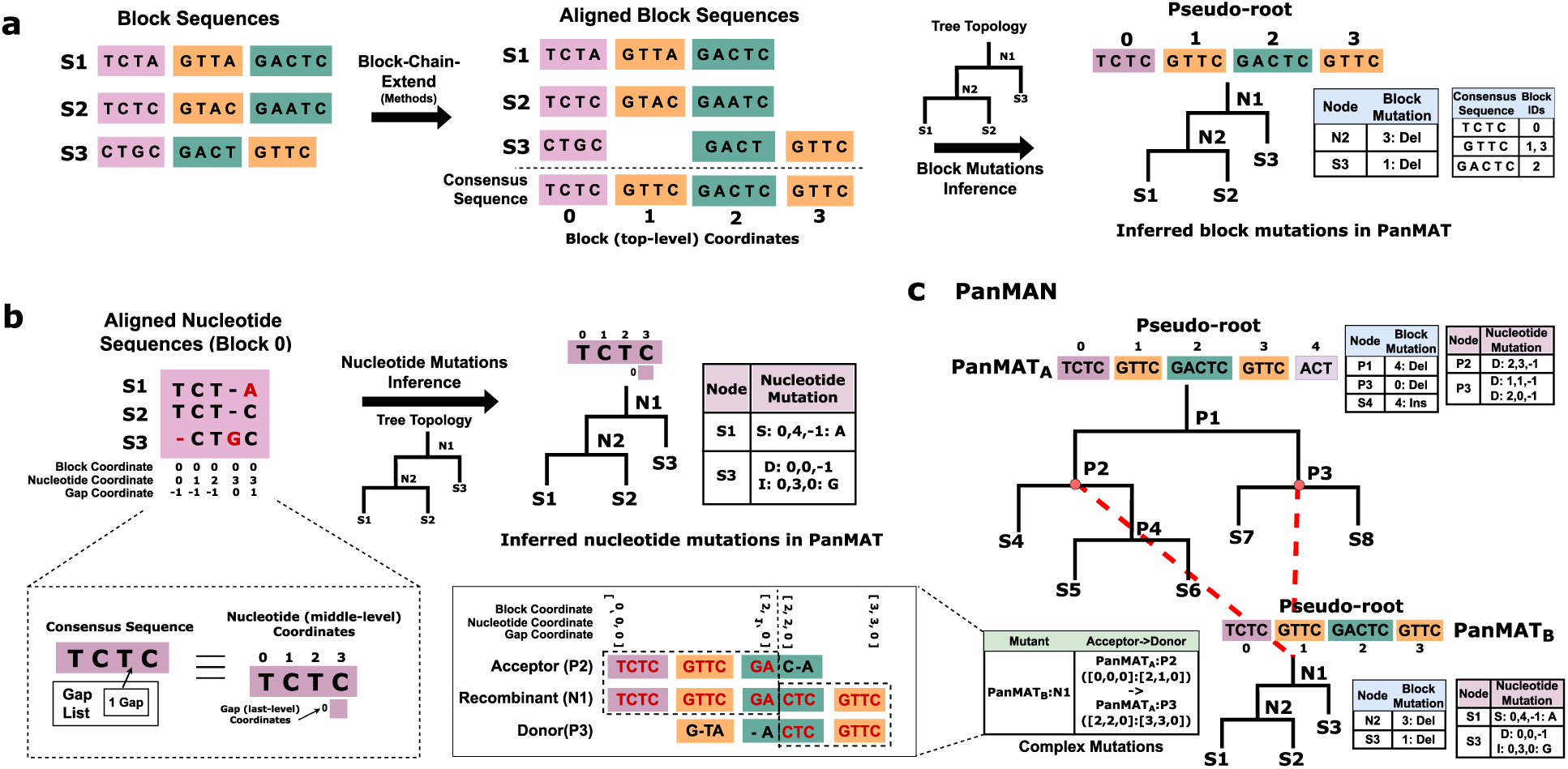
Overview of the PanMAN data structure. (**a**) Illustration of the top-level coordinate system in a PanMAT. The block sequences are first linearized with the help of multiple-sequence alignment and stored at the ‘pseudo-root’ of the PanMAT. (**b**) Illustration of the middle and lower-level coordinate system in PanMAT. These levels are based on a nucleotide-level multiple-sequence alignment of the consensus sequence with all sequences within a block. (c) Illustration of how PanMAN generalizes PanMATs to store complex mutations, namely, recombination and horizontal gene transfer, using network edges.

### PanMAN provides excellent compression for storing microbial pangenomes

We first evaluated the potential benefits of the evolutionary compression approach in PanMANs. We utilized sequence data from six diverse microbial species (SARS-CoV-2, respiratory syncytial virus (RSV), human immunodeficiency virus (HIV), *Mycobacterium Tuberculosis*, *Escherichia Coli*, and *Klebsiella pneumoniae*) and compared the file sizes with four standard variation-preserving pangenomic formats (GFA, VG, GBZ, and PanGraph) that provide lossless sequence encoding (Fig. 2, Methods). Our dataset selection spanned a wide range of genetic diversity, genome lengths, and sequence collection sizes. For each dataset, we built the PanMANs using a custom, parsimony-based pipeline that starts with alignments and phylogeny inferred by PanGraph (Methods).

PanMAN stands out as the most compressible format among variation-preserving pangenomic formats. PanMAN provides large compression ratios over all others: 19.5 to 680-fold over GFA, 6.1 to 221-fold over VG, 5.2 to 42-fold over GBZ, and 26.1 to 614-fold over PanGraph. Unsurprisingly, the gains of evolutionary compression tended to be the highest for SARS-CoV-2, which had the largest number of sequences in the collection and the lowest genetic diversity, as indicated by the average pairwise Mash distance^38^ shown in Fig. 3a. To test the scalability of file formats, we varied the number of sequences in the collection and found that the compression trend remained consistent (Fig. 3b). That is, PanMAN remained the most compressible format even for smaller collections and the gap between PanMAN and other formats widened when more sequences were added. Our results suggest that PanMAN scales nearly as well or better than other formats as the size of the sequence collection gets larger. These compression results are remarkable since PanMAN also has superior representative power.

**Figure 3:**
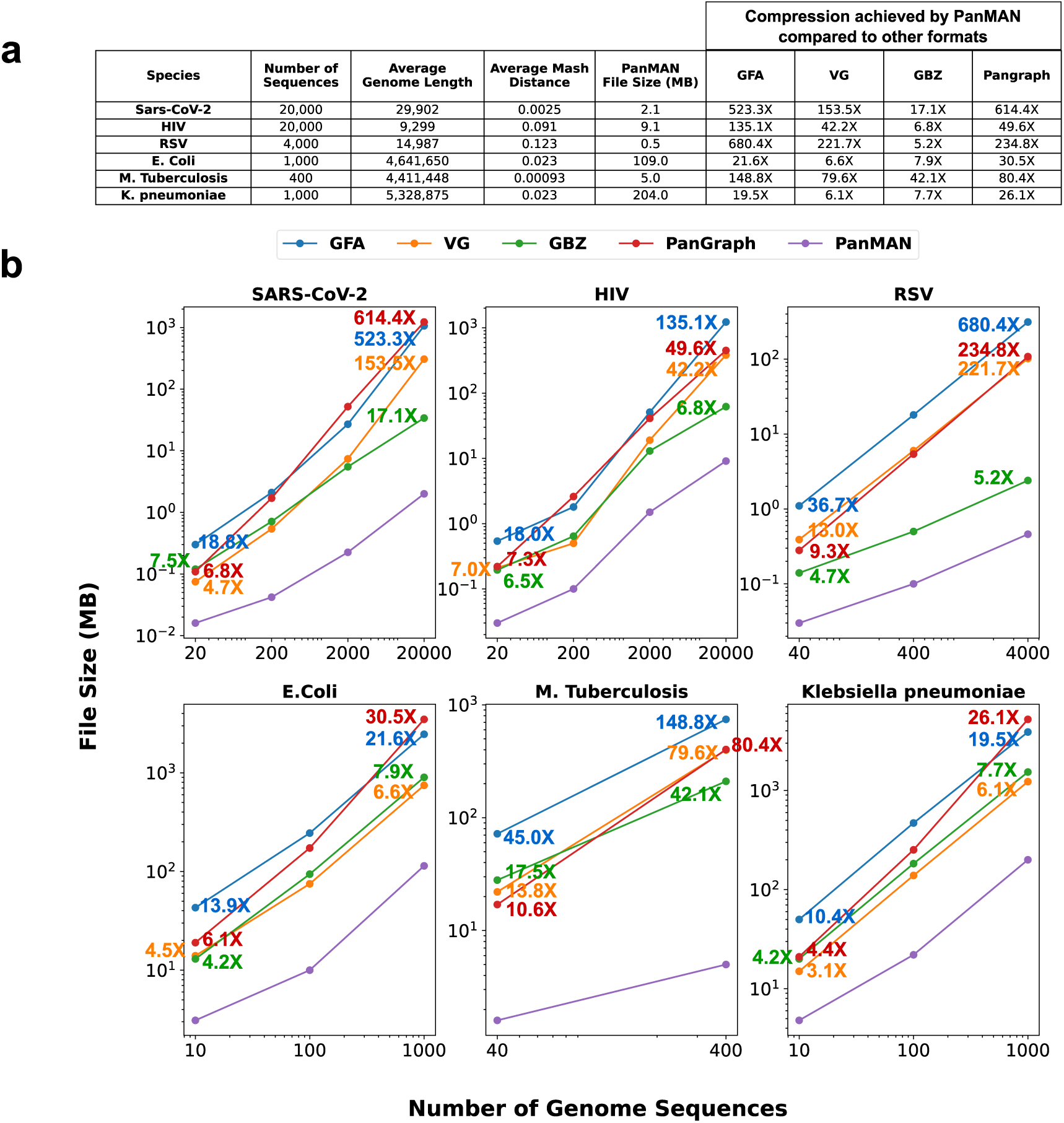
File sizes of PanMAN and other lossless pangenomic formats evaluated on six microbial datasets. (**a**) Compression achieved by PanMAN file format compared to other lossless pangenomic formats. (**b**) Scalability of PanMAN file format: compression achieved by PanMAN compared to other formats as the number of genome sequences increased in the collection. The numbers over markers indicate the file sizes relative to PanMAN. Both axes are on a logarithmic scale.

PanMAN is also competitive against file formats that do not explicitly represent the alignment or variation among sequences and are primarily designed for achieving high compression for similar genomes, namely MiniPhy^29^, AGC^28^, GZIP, ZIP, XZ, SAM^39^, BAM^39^, and CRAM^40^ (Supplementary Table 1). Most of these formats serve the sole purpose of data storage and retrieval. Notably, with the exception of MiniPhy, PanMAN outperforms all formats in storage efficiency on all datasets. Relative to MiniPhy, PanMAN achieves significantly higher compression rates (4.4 to 20-fold) for bacterial genomes and lower rates (0.6 to 0.9-fold) for viral genomes. This might be because encoding mutations relative to full sequences done in PanMANs is costlier for the considerably smaller viral genomes. However, the compression gap between MiniPhy and PanMAN for viral genomes also narrowed as the size of the datasets increased, suggesting that PanMAN scales better. Our results make a compelling case for PanMANs to be used as a standard data-sharing and storage format for large collections of related genomes.

### PanMAN improves the representative power of pangenomes

PanMAN has been crafted to represent a rich set of biologically meaningful information that current pangenome formats lack (Fig. 1). Some information in PanMAN is explicitly stored, such as mutations, phylogeny, annotations, and root sequence, whereas other information can be conveniently derived, such as ancestral sequences, multiple whole-genome alignment (m-WGA), and genetic variation. We have developed a set of algorithms in order to modify and extract useful information from PanMANs and packaged it under a software utility, called *panmanUtils*. Fig. 4 provides an overview of the different functionality that *panmanUtils* currently supports. Below, we describe how each piece of information is represented in a PanMAN or the algorithm using which it can be extracted in *panmanUtils*.

1. **Phylogeny.** At the center of a PanMAN is its corresponding phylogeny, which consists of single or multiple trees (PanMATs) connected by edges to form a phylogenetic network. Using *panmanUtils*, the topology of the phylogenetic network can be exported in two different ways: i) as a single network in the “extended Newick” format^41^ and ii) as multiple Newick files, one corresponding to each PanMAT topology, and an additional file describing the complex mutations resulting in network connections between the different PanMATs (Supplementary Fig. 1e).
2. **Multiple whole-genome alignment (m-WGA).** The coordinate system of PanMAT allows whole-genome alignment of all sequences in the collection, including inferred ancestral sequences, to be easily extracted through a single tree traversal (Fig. 4, Methods). Through *panmanUtils*, this m-WGA can be extracted for each PanMAT in a PanMAN in the form of a UCSC multiple alignment format (MAF). Specifically, each “alignment block” in an MAF corresponds to one homologous block inferred in the PanMAT, as shown in Fig. 4.
3. **Observed sequences and inferred ancestral genomes.** Any combination of sequences in a PanMAN, internal or those observed at tips, can be extracted by simply removing gaps from the m-WGA derived using the aforementioned method (Fig. 4).
4. **Genomic variation (VCF or GFA).** Since PanMAN explicitly stores mutations, variations are easily derivable. An interesting feature of *panmanUtils* is that it can also extract the variations of all sequences in a PanMAT with respect to an arbitrary reference sequence in the form of a Variant Call Format (VCF) file (Fig. 4, Methods). Moreover, to allow cross-compatibility with other formats, *panmanUtils* also supports the extraction of a GFA file representing the pangenome (Fig. 4, Methods). This allows PanMAN-extracted GFAs to be useful for read mapping and variant-calling applications by leveraging existing pangenomic tools (Supplementary Fig. 2).
5. **Annotations.** The utility also incorporates a function by which nodes of the PanMAN can be manually annotated with additional metadata, such as date, location, or lineage name, which can have various biological applications, such as genomic epidemiology (Fig. 4, Methods).

**Figure 4:**
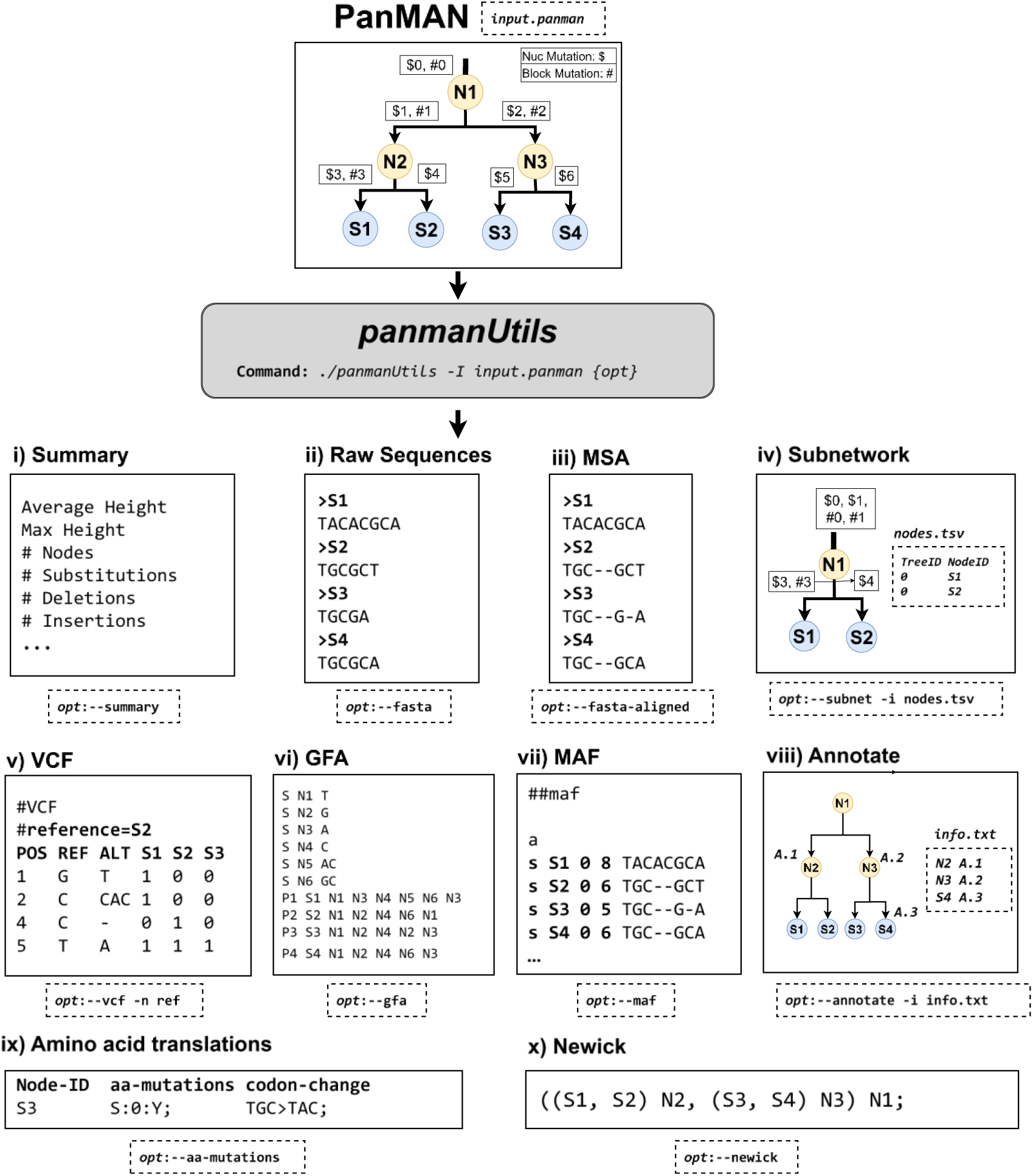
An overview of the different functionalities provided in *panmanUtils*. Starting from the input PanMAN (top), the panels show the following biologically relevant operations and their associated subcommands that can be performed using *panmanUtils*: (i) providing node and network-level statistics, (ii) extracting raw sequences at the tips or internal nodes in FASTA format, (iii) extracting the multiple-sequence alignment (MSA) of tip or internal sequences in FASTA format, (iv) extracting a PanMAN corresponding to a subnetwork in the input PanMAN based on the provided list of node identifiers in the tab-separated values (TSV) format, (v) extracting variation in VCF format present in all the sequences of a single PanMAT in a PanMAN with respect to a user-defined reference sequence (S2, in this case), (vi) converting any PanMAT in a PanMAN to a Graphical fragment assembly (GFA) file representing the pangenome (vii) extracting multiple whole genome alignment (m-WGA) for each PanMAT in a PanMAN in the form of multiple alignment format (MAF), (viii) annotating any node in the PanMAN with a custom string (e.g., lineage name) provided in a TSV format file, (ix) translating DNA mutation annotations to amino acid mutations with the reference being the root sequence, and (x) extracting multiple Newick files, one corresponding to each PanMAT topology, and an additional file describing the complex mutations resulting in network connections (if exist) between the different PanMATs.

The utility also incorporates several other useful functions, such as extracting subnetworks, providing summaries, rerooting PanMATs, or translating protein-coding sequences, which are described in greater detail in Methods. Most utility functions are parallelized using multi-threading and have reasonable runtime and memory requirements (Methods, Supplementary Tables 2 and 3).

### PanMANs support an arbitrary choice of algorithms for mutational and evolutionary inferences

By virtue of simultaneously storing phylogeny and mutations, PanMAN effectively represents ancestral characters. To this point in the manuscript, we only used parsimony for ancestral state reconstruction. Here, we demonstrate that PanMAN’s representation strategy can be applied to any inference method for inferring the phylogeny and ancestral states. Ancestral character reconstruction (ACR) is a well-established and active field of study, with several approaches based on parsimony, maximum likelihood, and Bayesian methods proposed in recent years^42–49^. To demonstrate the generalizability of PanMANs, we implemented a likelihood-based ACR approach, called Marginal Posterior Probabilities Approximation (MPPA)^47^, in *panmanUtils*, using which we inferred for different datasets the branch mutations on trees, which were constructed using a maximum likelihood-based tool called IQTree2^50^ (Methods). Compared to those obtained with the Fitch algorithm, the MPPA-based PanMANs were 2.2 to 7-fold larger in file sizes (Supplementary Fig. 3). An increase in file sizes is expected, as for any given alignment, the Fitch algorithm is guaranteed to annotate the fewest number of mutations on the branches of the tree but it may be less accurate than likelihood-based approaches for reconstructing ancestral sequences^42^. Despite larger file sizes, the MPPA-based PanMANs still provided 1.3 to 275-fold compression over other pangenome formats on the different datasets (Supplementary Fig. 3). Due to increased computational complexity, the MPPA-based PanMANs also took 3–6x longer to generate than Fitch-based PanMANs. Our experiments highlight that various inference methods offer different trade-offs between accuracy, compressibility, and computational complexity and PanMAN could serve as a valuable instrument for investigating these trade-offs in future studies.

### PanMAN unveils a detailed and accurate view of SARS-CoV-2 evolutionary and mutational history

The ability of a pangenomic data structure to simultaneously record mutational and evolutionary histories promises to have powerful applications in genomic epidemiology^8,51,52^, evolutionary biology^53,54^, and metagenomics^55^, among others. For instance, the comprehensive SARS-CoV-2 MATs produced by the UShER toolkit^30^ found widespread use during the COVID-19 pandemic, including naming variants^6,7^, studying local outbreaks^10,11^, predicting the fitness of different mutations^9^, and performing wastewater-based epidemiology^56^. However, several of these studies were also adversely impacted because of MAT’s inability to record indels and complex mutations^9,56,57^. For example, several studies have identified indels as key mutations defining many SARS-CoV-2 lineages and have found them to be associated with the enhanced fitness of the pathogen^57,58^. Moreover, a single tree representation that MAT uses for depicting SARS-CoV-2 evolution is inaccurate since the pathogen is known to have experienced a modest level of detectable recombination^59–61^.

We sought to explore if the PanMANs could address the above limitations of a UShER-MAT. We started by generating a PanMAT using *panmanUtils* for 20,000 SARS-CoV-2 sequences which spanned 1000 Pango lineages^6^. For a consistent comparison, we used the same tree topology as the UShER-MAT and the multiple sequence alignment derived from MAFFT^62^ (Methods). Specifically, mutations were annotated on the PanMAT using a parsimony-based algorithm of Fitch et al.^63^ (Methods). We also transferred the 1000 Pango lineage root annotations from the UShER-MAT to their corresponding locations in the PanMAT.

We found that PanMAN reconstructs similar single nucleotide mutation histories as UShER. Fig. 5a illustrates the number of substitutions, insertions, and deletions annotated in PanMAN and the bases affected in each. UShER’s MAT omits insertions and deletions (indels), which occur less frequently than substitutions but impact approximately four times more bases (Fig. 5a). Regarding substitutions, UShER-MAT and PanMAN show high concordance (Fig. 5b(i)), with PanMAN recovering over 99.7% of UShER-MAT substitutions. However, PanMAN records about 10% more substitutions on reference genome coordinates than UShER-MAT. This discrepancy arises primarily because UShER uses reference-based pairwise alignments, while PanMAN employs a multiple-sequence alignment (MSA) (Methods). PanMAN identifies 10,162 substitutions (9.9%) at MSA positions where the reference sequence (*Wuhan-Hu-1*, RefSeq: *NC_045512.2*) has no character, which UShER-MAT does not represent due to its reference-based system. High concordance is also observed between UShER-MAT and PanMAN when comparing substitutions from the reference to Pango lineage roots, with a Jaccard similarity of 0.968.

**Figure 5:**
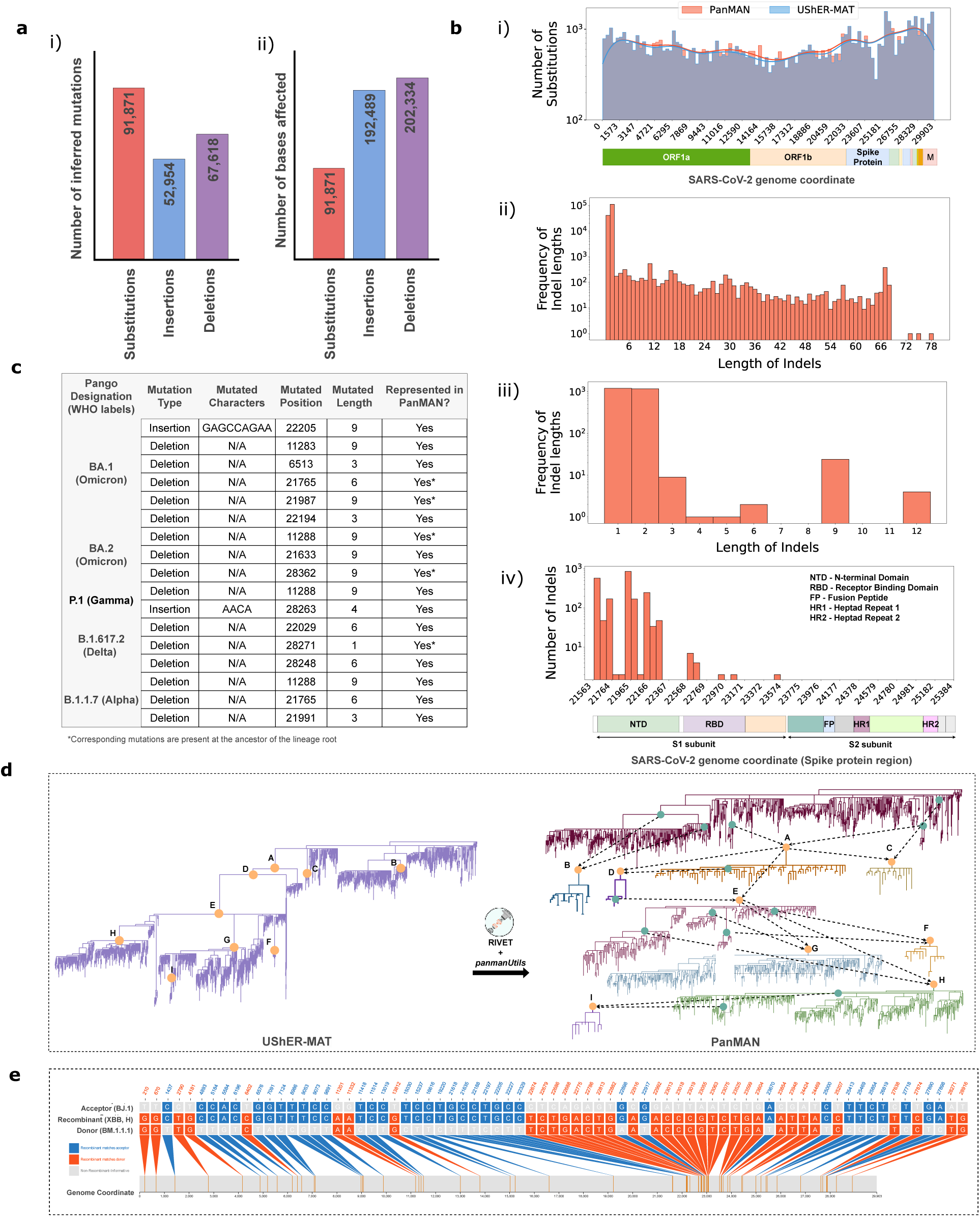
Exploration of SARS-CoV-2 mutational and evolutionary landscape on 20,000 sequences using PanMAN. (**a**) Illustrates the (i) number of inferred mutations and (ii) number of bases affected by inferred point mutations. (**b**) Landscape of mutations inferred in PanMAN. (i) The distribution of inferred substitution in PanMAN and UShER-MAT across SARS-CoV-2 reference genome coordinates, (ii) The frequency of inferred indel lengths across SARS-CoV-2 reference genome coordinates, (iii) the frequency of inferred indel lengths in the spike protein region, and (iv) the distributions of inferred indels in the spike protein region. (**c**) PanMAN accurately infers the lineage-defining indels, reported in a previous study^57^ for different variants of concern (VOCs). (**d**) PanMAN represents a comprehensive view of the SARS-CoV-2 evolution through recombination. The recombination events were inferred using RIVET and the *panmanUtils* splits the single-tree PanMAN into a network of trees. (**e**) Recombination-informative sites (that is, sites where the recombinant node matches either but not both parent nodes) are shown for XBB. The figure was generated using RIVET.

Our SARS-CoV-2 PanMAN offers a detailed representation of the evolution of indels. Genome-wide, indel frequency decreases with length (Fig. 5b(ii)). This pattern persists in the Spike protein, where indels of lengths in multiples of three are notably more common, indicating strong purifying selection against frame-shifting mutations in the Spike protein compared to the rest of the genome (Fig. 5b(iii)). Indels longer than 12 bases are extremely rare in the Spike protein. To verify the accuracy of indel inference, we compared our results to a previous study^57^ on lineage-defining mutations for variants of concern (Omicron sub-lineages BA.1 and BA.2, Delta B.1.617.2, Gamma P.1, and Alpha B.1.1.7) and found that all reported indels were correctly annotated in PanMAN (Fig. 5c). Within the Spike protein, indels are very rare in the S2 subunit (Fig. 5b(iv)) that plays an essential role in viral fusion and entry into host cells^64^. Spike protein indels are most frequent in the N-terminal domain (NTD) – one of the most genetically modified regions of the Spike protein – of the S1 subunit. This observation is concordant with the findings of a previous study^65^, as NTD indels are thought to be an important mechanism for immune evasion and adaptive evolution of SARS-CoV-2^66^.

Further, to account for recombination, we inferred recombination events in the UShER-MAT using the RIVET pipeline^59,67^ and using *panmanUtils*, split the PanMAT based on recombination events into multiple interconnected PanMATs forming the final PanMAN (Fig. 5d, Methods). The remaining mutations were not changed during the splitting. The final PanMAN has roughly the same file size as the original PanMAT but offers a more accurate and comprehensive view of the SARS-CoV-2 evolution (Fig. 5d). Notably, several recombinants that are designated under the Pango nomenclature, like XBB, were accurately indicated on the PanMAN (Fig. 5e).

Overall, our analysis provides a compelling case study on PanMAN’s ability to offer a detailed and accurate view of the evolutionary and mutational histories of different species.

## Discussion

We presented a novel data representation, PanMAN, that provides excellent compression to store a collection of evolutionarily related genome sequences and greatly improves on the representative power of pangenomes. Our work has laid a strong foundation for further enhancing the scalability, accuracy, generalizability, and accessibility of PanMANs and its associated tools.

Although PanMAN is a compact and scalable format, constructing them is computationally intensive due to its reliance on existing tools (such as PanGraph) for phylogenetic and m-WGA construction (Supplementary Table 4). In our future work, we plan to enable the construction of larger PanMANs consisting of hundreds of thousands to millions of sequences with the help of parallel algorithms and GPU acceleration^68^. Besides scalability, visualization is another important aspect of pangenomic analysis. While graph genomes have associated visualization tools^69–72^, and PanMAN can be converted to these structures via *panmanUtils*, graph genomes are quite challenging to interpret and analyze. This is another aspect where PanMANs could prove valuable – our earlier work on visualizing UShER-MATs using Taxonium and the Treenome Browser^73,74^ could be easily adapted to PanMANs in the future. In addition, we plan to develop read mapping tools better optimized for PanMANs, however, as we demonstrated in this manuscript, PanMANs have compatibility with existing pangenomic tools that overcome reference bias. Lastly, PanMANs have so far only been demonstrated on viruses and prokaryotic genomes. While many ideas in PanMANs are generalizable, we also foresee challenges when applying them to eukaryotic species that involve multiple chromosomes, reproduce sexually, and undergo frequent recombination. Our future work will also entail addressing these challenges.

The accuracy of PanMANs hinges on the accuracy of inference algorithms. PanMAN currently relies on existing approaches, such as PanGraph and the Fitch algorithm, for phylogeny, alignment, and ancestral character reconstruction. However, we have also demonstrated that PanMAN is versatile and can work with any choice of underlying inference algorithm. These areas are a focus of ongoing research^42,47,50,75–80^ and PanMAN might also serve as a useful platform to support these studies. Likewise, recombination and HGT inference methods are also active areas of research^59,60,81–83^ and will benefit from PanMANs for qualitative analysis.

Overall, our work has the potential to transform pangenomic analysis. This is because PanMANs are not just efficient, they also provide a unique mechanism to unify a variety of biologically meaningful information, such as phylogeny, mutational history, genomic variation, and m-WGA, under a common format. This has far-reaching applications in epidemiological, microbiological, metagenomic, ecological, and evolutionary studies.

## Methods

### Datasets

We evaluated pangenomic file formats on six datasets consisting of 1) 20,000 SARS-CoV-2, 2) 1,000 *Escherichia Coli*, 3) 20,000 Human Immunodeficiency Virus (HIV), 4) 4,000 Respiratory Syncytial Virus (RSV) 5) 400 *Mycobacterium Tuberculosis*, and 6) 1,000 *Klebsiella pneumoniae* genome sequences. The SARS-CoV-2 genome sequences were collected from GISAID^4^ online database, the HIV sequences were collected from a database maintained by Los Alamos National Laboratory (LANL)^84^, and the remaining were downloaded from NCBI RefSeq^85^. The details of these pangenomic datasets, including NCBI and GISAID IDs, are provided in Supplementary Tables. GISAID maintains more than 15 million SARS-CoV-2 genome sequences with over 3000 Pango lineages. For our purposes, we selected the 1,000 largest SARS-CoV-2 lineages and downsampled the dataset to 20 random samples per lineage to get 20,000 sequences. *Escherichia Coli*, RSV, *Mycobacterium Tuberculosis*, and *Klebsiella pneumoniae* genome sequences were randomly sampled from the collected sequences from the NCBI database; details can be found in Supplementary Tables. For HIV, we removed the 5853 problematic sequences flagged by LANL and randomly sampled 20,000 sequences from the remaining 20,525 sequences.

### PanMAN data structure: detailed description

A PanMAN can be composed of a single or multiple PanMATs. We first describe the PanMAT structure below, followed by a discussion on how they can be combined to form a PanMAN.

A PanMAT is designed to represent a collection of sequences that have evolved solely through vertical descent, i.e., wherein each sequence in the collection has mutated from a single ancestral sequence. PanMAT maintains a phylogenetic tree in which the sequences in the collection are represented at tips (Fig. 2). However, instead of storing individual sequences in the collection, PanMAT maintains a single sequence at the root of the tree, with branches of the tree annotated with mutations. As a result, any tip sequence can be derived from the root sequence by applying the series of mutations along the path from the root to that tip. Because many mutations are shared and can be compactly stored relative to the complete sequences, this method provides large gains in storage efficiency. This idea was introduced in UShER as “evolutionary compression”^30^. Compared to UShER’s MAT, PanMAT introduces a reference-free three-level coordinate system that is inspired by PanGraph^22^ to accommodate indels and structural rearrangements.

At the top level, PanMAT’s coordinate system is denoted by *blocks* (Fig. 2a), which represent homologous segments between sequences. In other words, if the same block is present in two sequences, it means that the segments represented by the block in the two sequences could be aligned and are inferred to share an ancestry. Blocks may also signify segments unique to a particular sequence, lacking homologs in other sequences. Multiple blocks in a PanMAT are linearized using the *block-chain-extend* algorithm (described in a subsequent subsection) and each block is assigned a unique identifier based on its position in this linear sequence (Fig. 2a). This block sequence is stored at the ‘pseudo root’, which acts like a parent to the actual phylogenetic root in the PanMAT (Fig. a). Each block also stores a ‘consensus sequence’ which is derived from the alignment of sequences in the collection (Fig 2b). Blocks can be inserted or deleted on branches of PanMAT, and they determine the exact sequence of blocks present at any node of the tree. This allows a convenient mechanism to represent large indel mutations in a PanMAT. While blocks that are duplicated or rearranged are assigned unique identifiers in PanMAT’s linear coordinate system, PanMAT separately maintains lists of the blocks that are homologs (Fig. 2a), and homologous blocks share a single consensus sequence.

The lower two levels of the PanMAT’s coordinate system are used to represent the specific coordinate within the block in the presence of small indels. This system is based on the multiple sequence alignment (MSA) of the consensus sequence with other sequences in the collection (Fig. 2b). Since in the MSA, the consensus sequence is interspersed with gaps, the middle level of the PanMAT’s coordinate system corresponds to the position of the last consensus nucleotide and the last level corresponds to the position of the gap since that nucleotide (Fig. 2b). The lengths of the gaps following every consensus nucleotide are maintained at the pseudo root.

PanMAN provides a generalization of PanMATs to also store complex mutations, namely recombination and horizontal gene transfer (HGT), which violate the vertical inheritance assumption of PanMAT as with these mutations, a sequence can inherit genetic information from two or more parent sequences. PanMAN uses network edges to connect two nodes in the same or different PanMATs to the root of a new PanMAT. These network edges store the complex mutation that results in the new PanMAT’s root sequence. For recombination, the network edges store the list of breakpoint coordinates in the two source node sequences where the recombination took place (Fig. 2c). For HGT, the network edges store the start and end coordinates of the gene in one sequence, and the coordinate in the second sequence at which the gene is inserted (Fig. 2c).

### PanMAN’s file format

PanMAN utilizes oogle’s protocol buffer (protobuf, https://protobuf.dev/), a binary serialization file format, to compactly store PanMAN’s data structure in a file. Supplementary Fig. 4 provides the .proto file defining the PanMAN’s structure. At the top level, the file format of PanMANs encodes a list (declared as a *repeated* identifier in the .protof file) of PanMATs. Each PanMAT object stores the following data elements: (a) a unique identifier, (b) a phylogenetic tree stored as a *string* in Newick format, (c) a list of node objects, ordered according to the pre-order traversal of the tree topology, (d) a block mapping object to record homologous segments identified as duplications and rearrangements, which are mapped against their common consensus sequence; the block-mapping object is also used to derive the pseudo-root, e) a gap list to store the position and length of gaps corresponding to each block’s consensus sequence. Each node object encodes a) a list annotations field (*string*), which allows any node to be annotated with a custom string and later searched by these annotations, and b) a list of mutations object. Each mutation object encodes the node’s block and nucleotide mutations that are inferred on the branches leading to that node. If a block mutation exists at a position described by the Block-ID field (*int32*), the block mutation field (*bool*) is set to 1, otherwise set to 0, and its type is stored as a substitution to and from a gap in Block mutation type field (*bool*), encoded as 0 or 1, respectively. In PanMAN, each nucleotide mutation within a block inferred on a branch has four pieces of information, i.e., position (middle coordinate), gap position (last coordinate), mutation type, and mutated characters. To reduce redundancy in the file, consecutive mutations of the same type are packed together and stored as a mutation info (*int32*) field, where mutation type, mutation length, and mutated characters use 3, 5, and 24 bits, respectively. PanMAN stores each character using one-hot encoding, hence, one “Nucleotide Mutations” object can store up to 6 consecutive mutations of the same type. PanMAN’s file also stores the complex mutation object to encode the type of complex mutation and its metadata such as PanMATs’ and nodes’ identifiers, breakpoint coordinates, etc. The entire file is then compressed using XZ (https://github.com/tukaani-project/xz) to enhance storage efficiency.

### PanMAN construction algorithms in *panmanUtils*

*panmanUtils* includes multiple algorithms to construct PanMANs in addition to supporting various functionalities to modify and extract useful information from PanMANs. In this section, we delve into the construction algorithms in *panmanUtils*. Each of these algorithms plays a crucial role in enhancing the toolkit’s functionality and enabling comprehensive pangenomic analyses. *panmanUtils* is implemented in C++, built, and compiled using CMake v3.16.3 and g++-10.3, respectively. It is a highly optimized program implemented to parallelize across all available virtual cores on a CPU using Intel’s Threading Building Blocks^86^ (TBB v2019) library. It utilizes the Boost C++ library^87^ v1.17.0 to provide the program options via the command line and protobuf v3.20.3 to compactly store PanMAN files.

### PanMAN construction pipeline

PanMAN can be composed of a single or multiple PanMATs. To construct a PanMAN, we start with a PanMAN consisting of a single tree (or PanMAT) and then split it up into a network of multiple PanMATs using the inferred complex mutations provided as input (Supplementary Fig. 5).

To construct the starting single-tree PanMAN representing a collection of sequences, *panmanUtils* require two inputs a) tree topology representing the phylogenetic relationship of the input sequences and b) a pangenomic data structure (PanGraph or GFA or FASTA) representing the multiple-sequence alignment (MSA) corresponding to the sequence collection. We built the inputs as described in the subsequent subsection.

If PanMAN is constructed from a graph-based pangenomic data structure (PanGraph or GFA), the initial block sequence at the tip of the tree is known from the paths in the input, where pancontigs (nodes) in PanGraph (GFA) data structure are translated as blocks in PanMAN (Supplementary Fig. 5a). It is critical to recognize that a few microbial species, such as E. coli and Klebsiella pneumonia, have circular genomes, which implies the absence of a defined start and end point and can lead to sequence misalignment within a pangenome, erroneous mutation inference, and inefficient PanMANs. To address this, we employ a rotation algorithm (described in the next subsection) to align tip sequences better (Supplementary Fig. 5a). Next, a sequence of blocks is generated using a block-chain-extend algorithm to linearize block sequences and handle duplications and rearrangements in the input graph pangenome (Methods), where the block at each position in the sequence is assigned a unique identifier (top-level coordinate). This block sequence is stored at the ‘pseudo root’, which acts like a parent to the actual phylogenetic root in the PanMAN (Supplementary Fig. 5a). Next, an ancestral character reconstruction (ACR) methodology is used to infer block sequences at the internal nodes of the tree and subsequently, inferring block mutations on the branches (Supplementary Fig. 5a) by simply finding differences with the parents’ block sequence.

PanGraph also provides variations within each block through an MSA, which is directly translated to the second and third-level coordinates of PanMAN’s coordinate system. Again, an ACR algorithm is applied to each aligned nucleotide coordinate and infers nucleotide mutations on the branches (Supplementary Fig. 5a). By default, *panmanUtils* uses the Fitch algorithm (parsimony-based) for ACR, however, an arbitrary methodology can be adopted in *panmanUtils* (ACR methods). Since the GFA format does not encode variations within a node, PanMANs built from a GFA do not encode inferred nucleotide mutations (Supplementary Fig. 5a). In the case of an MSA as the input, *panmanUtils* treats the whole alignment as one block that is present in every sequence and infers only the nucleotide mutations within this block on the branches (Supplementary Fig. 5a). Therefore, as expected, we observed that *panmanUtils* constructs the most memory-efficient PanMANs using a PanGraph, as it leverages the evolutionary compression at both block and nucleotide levels.

Finally, to construct a network of trees in PanMANs to represent a complex mutation, namely recombination and HGT, *panmanUtils* requires two inputs a) a list of PanMANs constructed in the previous step and b) metadata of the complex mutations (Supplementary Fig. 5b). The metadata includes, for recombination, a list of breakpoint coordinates in the two source node sequences where the recombination took place, or for HGT, the start and end coordinates of the gene in one sequence, and the coordinate in the second sequence at which the gene is transferred. If both the parents and child nodes belong to the same tree in the PanMAN, *panmanUtils* breaks the tree into two trees, assigning a unique identifier to each tree, such that the source node belongs to the first tree and the child node is at the root of the second tree (Supplementary Fig. 5b). If parents and child nodes belong to multiple trees and the child node is not at the root of the tree, it is re-rooted such that the child node becomes the root of this tree. Finally, PanMAN stores the complex mutation using the four coordinates, the nodes’ identifiers, and the list of the tree indices containing these nodes.

### Rotation algorithm to deal with circular sequences

A few microbial species, such as *E. coli* and *Klebsiella pneumonia*, have circular genomes forming a closed loop, which implies the absence of a defined start and end point. In the genome assemblies of these species, the start positions are typically chosen arbitrarily. Because PanMAN’s coordinate system expects sequences to be linearized, sequences with different starting points can result in unnecessary duplication of blocks, leading to storage-inefficient PanMANs. To address this issue, we employ a variant of the Needleman-Wunsch algorithm, tailored for rotating sequences to a common starting point (Supplementary Fig. 5a). In this algorithm, the scoring for cell *C*(*i*,0) depends on *C*(*i*-1,0) and two cells in the last row of the traceback matrix, i.e., *C*(*i*,N) and *C*(*i*−1,N). Therefore, accommodating the circular nature of these genomes and generating precise alignment and rotation offset. We arbitrarily choose a reference sequence and align the rest of the sequences to find the optimal rotational offset. However, aligning raw nucleotide sequences directly, as also proposed in *Rotate*^88^, is computationally demanding. To mitigate this, we leverage block sequences, which are substantially shorter, to find the optimal rotational offset—significantly reducing computational load. Once these offsets are determined, the block sequences are rotated accordingly, and these rotational values are incorporated into the PanMAN file format, to reproduce the correct block order required by other tools in the *panmanUtils*, such as for raw sequence extraction.

### Block-Chain-Extend Algorithm

In PanMAN, block position acts as the block identifier, and block mutation inference occurs at each position separately. Hence, the block sequences need to be perfectly aligned, which means that the relative position of any pair of blocks needs to be the same in every sequence in the pangenome. However, duplications and rearrangements violate this property. Therefore, we developed block-chain-extend, which is inspired by the seed-filter-extend algorithm, to handle rearranged and duplicate blocks in input graphs, by *renaming* them while preserving the homology information. Unlike the seed-filter-extend algorithm, here a reference chain of blocks is maintained to which every other block sequence is mapped, to create an alignment. First, the reference chain is chosen to be a block sequence of any arbitrary tip of the phylogenetic tree. Then, progressively for every other tip (or sample), the blocks in the reference chain that are homologous to the sample’s block sequence are marked in the table (Supplementary Fig. 5a). Next, the most optimal chain of marked cells from the table is found by maximizing the alignment score. Note that the cells in this chain are such that each of their coordinates is strictly increasing. The block identifiers in the sample are renamed to be the identifiers of the blocks in the reference chain that they align to and the blocks in the sample chain that do not align are added to the reference chain in their corresponding positions (Supplementary Fig. 5a). However, renaming the block’s identifier can lead to a loss of homology information, therefore, PanMAN maintains a one-to-many mapping of a consensus sequence to a list of block identifiers that are homologous (Supplementary Fig. 5a), such that biologically relevant information such as rearrangements and duplications is maintained.

### Ancestral Character Reconstruction (ACR) methods

We implemented two ACR methods in *panmanUtils*, namely, Fitch (parsimony-based) and MPPA (likelihood-based) to infer branch mutations.

Given aligned sequences of the tips of the tree, either nucleotide or block, the Fitch algorithm finds the most parsimonious state for every node of the tree (Supplementary Fig. 1c). Since multiple states may be most parsimonious at a node (Supplementary Fig. 1c), *panmanUtils* assigns any one of the most parsimonious states with preference given to consensus sequence base in case of nucleotide sequences and block in pseudo-root in case of block sequences. *panmanUtils* also allows ambiguous nucleotide bases in tip sequences using the IUPAC format (https://www.bioinformatics.org/sms/iupac.html), which are also resolved to a unique base using the above-mentioned strategy after inferring the most parsimonious nucleotide(s). When a branch leading to a node is found to carry a mutation, that is, the state assigned to the node differs from its parent, the inferred mutation (Supplementary Fig. 1c) is added to a list of mutations corresponding to the branches leading to that node.

MPPA, a likelihood-based ACR method, also used in the PastML^47^ software, is inspired by the Fitch algorithm. state frequencies by performing Felsenstein’s pruning algorithm^89^ and the bounded variant of the limited memory Broyden–Fletcher–Goldfarb–Shanno optimization algorithm^90^. These parameters are then used to compute the approximate marginal posterior probabilities of each state at each node. This is done by first performing a post-order traversal on the input tree (estimated using maximum-likelihood methods, such as Felsenstein’s pruning algorithm) to account for the information from each node’s descendants, followed by a pre-order traversal when this information is combined with information from other parts of the tree to compute the marginal probabilities. Once the marginal probabilities are computed, a subset of the states is selected for each node using a dynamic thresholding logic. Since the MPPA algorithm allows for uncertain (multiple) states at the internal nodes, we use IUPAC nucleotide codes to represent them. We note that *panmanUtils* only uses MPPA to compute nucleotide mutations and block mutations are still computed using the parsimony-based methods.

## Functionalities in *panmanUtils*

We now describe *panmanUtils* multiple algorithms to support various functionalities to modify and extract useful information from a PanMAN (*input.panman*).

### Subnetwork extract

*panmanUtils* supports the extraction of subnetworks from a network of trees in a PanMAN (*data.panman*) based on the provided list of node identifiers and corresponding tree identifiers in a TSV format text file (*nodes.tsv*) using the command: “*./panmanUtils -I data.panman --subnet nodes.tsv*”. *panmanUtils* implements this feature by traversing from the nodes listed in the file to the root and marking nodes along the way (Supplementary Fig. 1b). Since this traversal from each node can be executed independently, *panmanUtils* exploit multiple CPU threads to parallelize the algorithm, each thread handling one node at a time. Finally, the subnetwork consisting of marked nodes is extracted and the redundant nodes - those having only one child - are combined with their children until the network contains no redundant nodes (Supplementary Fig. 1b). Note that, if the requested nodes break the network structure in the PanMAN, *panmanUtils* implicitly keeps the nodes to maintain the hierarchy.

### Tip/internal node sequences and multiple sequence alignment (MSA) extract

*panmanUtils* supports extraction of raw (tip) and internal node sequences from a PanMAN in a FASTA format using the command: “*./panmanUtils -I data.panman --fasta*”. The implementation of this feature involves starting with the consensus sequence of each block at the pseudo-root of the PanMAN and performing a post-order traversal while applying the mutations along the way. Once a desired node of a tree is reached, the sequence representing the node has been computed and can be printed to the file. When traversing up from a node to its parent, the mutations on that branch also need to be reversed.

*panmanUtils* also supports extraction of multiple sequence alignment (MSA) of sequences for each PanMAT in a PanMAN in a FASTA format using the command: “*./panmanUtils -I data.panman --fasta-aligned*”. Since PanMAT’s coordinate system allows implicit storage of alignment of sequences, implementation of this feature in *panmanUtils* is similar to the above-mentioned algorithm, except for only printing the nucleotides that are present in each sequence, the gaps are also printed.

### Multiple Whole Genome Alignment (m-WGA) extract

*panmanUtils* supports extraction of m-WGA for each PanMAT in a PanMAN in the form of a UCSC multiple alignment format (MAF, https://genome.ucsc.edu/FAQ/FAQformat.html) using the command: “*./panmanUtils -I data.panman --maf*”. Similar to MSA extract, this functionality implementation requires a single traversal over each PanMAT to extract tip sequences. Next, in order to print m-WGA in the form of MAF, the start and end index of each block are computed for each sequence to break the alignment and print them, along with the start indexes and lengths of each block.

### Variant Call Format (VCF) extract

*panmanUtils* supports the extraction of variations of all sequences from any PanMAT in a PanMAN in the form of a VCF file (Supplementary Fig. 1a). A unique feature of *panmanUtils* is that it allows extraction of variations with respect to *any* reference sequence (*ref*) in the PanMAT, using the command: “*./panmanUtils -I data.panman --vcf -n ref*”. The variations of each sequence with respect to the reference sequence are simply computed by comparing each position of the alignment of the two sequences which is obtained from the MSA.

### Graphical fragment assembly (GFA) extract

*panmanUtils* supports converting any PanMAT in a PanMAN to a Graphical fragment assembly (GFA) file representing the pangenome (Supplementary Fig. 1d), using the command: “*./panmanUtils -I data.panman --gfa*”. In order to implement this algorithm, *panmanUtils* uses PanMAT’s block-wise alignment, where GFA nodes are identified as contiguous segments in each block. Next, the nodes that appear after one another are linked with edges and the nodes appearing in multiple sequences at the same positions in the alignment sequences are reused (Supplementary Fig. 1d). We further compress the generated graph by iteratively merging two nodes (Supplementary Fig. 1d), iff, two blocks occur right after each other in all the sequences. Finally, the nodes and edges of this graph along with the paths are printed in a GFA format file.

In order to determine if GFAs generated using *panmanUtils* from PanMANs could compete with standard pipelines in terms of graph complexity and read mapping accuracy and speed, the PanMANs constructed from the PanGraph alignments were converted into corresponding GFAs using *panmanUtils* and compared against the GFAs produced by the pangenome graph builder (PGGB) pipeline for the same sequence collections (described in a subsequent subsection).

### Other functionalities

*panmanUtils* also supports features such as summary extract, annotations, rerooting, and protein translation extract. The summary of a PanMAN can be extracted using the command: “*./panmanUtils -I data.panman --summary*”, which contains a summary of its geometric and parsimony information. This feature’s computation is parallelized using the parallel reduction paradigm. Any node in a PanMAN can be annotated with a custom string, stored in the annotations field (*string*) in the PanMAN’s protobuf file format, and later searched by these annotations, using the command: “*./panmanUtils -I data.panman --annotate info.txt*”, where *info.txt* is a tab-delimited file in which each line corresponds to a unique annotation. *panmanUtils rerooting* functionality allows the rerooting of any PanMAT in a PanMAN to a tip node in that PanMAT using the command:”*./panmanUtils -I data.panman --reroot -n tip-node*”. Finally, a protein translation file can also be obtained which contains the protein mutations between any two reference-based coordinates with the reference being the root sequence.

## Construction of baseline file formats

We compared PanMAN with four lossless variation-preserving pangenomic file formats (GFA, VG, GBZ, and PanGraph) for each analyzed dataset. Specifically, we generated GFA files using the Pangenome Graph Builder (PGGB) *v0.6.0*, configured to its default parameters, using the command: *./pggb -i data.fa -t 32 -n num_sequences -o output_dir*. The PGGB algorithm accepts raw genome sequences in FASTA format (*data.fa*) to assemble a comprehensive pangenome represented in GFA format (*data.gfa*). We used these generated GFA files from PGGB to construct pangenome represented in V and BZ files using “*vg convert”* and “*vg gbwt”* tools from the vg-toolkit *v1.56.0*, with the default parameters, using the commands: *./vg convert -p data.gfa -t 32 -v > data.vg* and *./vg gbwt -G data.gfa --num-threads 32 --gbz-format -g data.gbz*, respectively. The pangenome represented as PanGraph alignments are constructed from the raw sequences using the PanGraph tool *v0.7.3.* Apart from graph-pangenome, PanGraph also generates a guide tree built using the Neighbour-joining^91^ algorithm, a distance-based phylogenetic tree construction method, representing the phylogenetic relationship between the set of sequences in the pangenome. The PanGraph tool uses a pseudo-energy function, expressed as *E*=−*l*+*αNc*+*βNm*, to gauge the complexity of the resultant graph. Here, *l* represents the length of alignment that facilitates graph merging, *Nc* indicates the number of additional blocks formed, and *Nm* reflects the number of variations in the alignment used for merging. The user-defined parameters *α* and *β* modulate the degree of graph fragmentation and the intra-block variation, respectively. We optimized *α* and *β* for the datasets analyzed and used the command: *./pangraph build -k mmseqs -a α -b β data.fa | ./pangraph polish*, with other parameters set to default, to create the most memory-efficient PanGraph alignments (*data.json*) and guide tree in Newick format (*data.nwk*).

We also explored compression techniques (GZIP, ZIP, XZ, SAM, BAM, CRAM, AGC, and MiniPhy) and compared their file sizes with PanMAN files, for each analyzed dataset. In particular, we generated the GZIP compressed files (*data.fa.gzip*) from the raw genome sequences in FASTA format (*data.fa*) using the command: “*gzip data.fa*” with Z P v . . Similarly, Z P-compressed and XZ-compressed files are generated using the command: “*zip -r data.fa.zip data.fa*” with Z P v . and “*xz data.fa*” using XZ utils v , respectively. We used minimap ^92^ software, v2.28, to produce SAM files (*data.sam*). To do so, we randomly sampled one genome sequence from *data.fa* as the reference sequence (*ref.fa*) and used the command: “*minimap2 -a -t 32 ref.fa data.fa*”, with other parameters set to default. These SAM files were converted to BAM files (*data.bam*) with SAMtools^39^, using the command: “*samtools view -S -b data.sam*”. Finally, we used SAMtools and the reference genome sequence to convert previously generated BAM files to CRAM files (*data.cram*) using the command: “*samtools view -C -T ref.fa data.bam*”. We set all the parameters to default while using the samtools. We built AGC files from the raw sequence with the AGC tool v3.1 using the command: *./agc create data.fa > data.agc*, with default settings. Finally, we used MiniPhy v0.4.0^29^ to generate MiniPhy-XZ compressed files, with default settings.

### Read mapping on GFAs

We produced GFAs by converting PanMANs (*data.panman*) constructed from the PanGraph alignments using *panmanUtils*, using the command: *./panmanUtils -I data.panman --gfa* and compared against the GFAs produced with PGGB (described in the previous subsection). To determine the read mapping accuracy and speed on GFAs, we first simulated reads from one arbitrary haplotype from three microbial datasets (SARS-CoV-2, RSV, and HIV) using the ART sequencing read simulator^93^, using the ‘HiSeqX TruSeq’ sequencing system (a read length of 150 bases and coverage of 60X). Then, we mapped these simulated reads on GFAs using an existing read-mapping tool called Giraffe^94^, and utilized the reported number of reads mapped (Supplementary Fig. 2).

### Constructing likelihood-based phylogenetic trees

To infer mutations using the MPPA algorithm, we used likelihood-based phylogenetic trees and employed the F81 evolutionary model^89^, where each base frequency was calculated by taking a fraction of the base count and the total number of sites in the alignment. We constructed the likelihood-based trees in two steps. First, the generated PanMANs (*data.panman*) constructed from the PanGraph alignments are used as input to *panmanUtils* to build the MSA (*data.msa*) in the FASTA format using the command: *./panmanUtils -I data.panman --msa*. Then IQTree2 *v2.3.2* was used to construct the likelihood-based tree (*data.iqtree.nwk*) using the command: *./iqtree2 - s data.msa -T 32*, with default settings.

## SARS-CoV-2 mutational and evolutionary landscape

We downsampled the 16 million SARS-CoV-2 sequences maintained by GISAID to 20,000 sequences (*sars_20000.fa*), spanning the 20 samples per top-1000 Pango lineages. We further downsampled the comprehensive SARS-CoV-2 UShER-MAT maintained by UCSC^95^ to the desired 20,000 SARS-CoV-2 sequences (*sars_20000.mat.pb*) to extract inferred lineage-defining substitutions in UShER-MAT. However, the UShER-MAT maintained by UCSC masks many problematic sites^96^, and to recover all of them, we reconstructed UShER-MAT maintaining the same tree topology. Specifically, the tree topology was first extracted using *matUtils*^95^, using the command: “*matUtils extract -i sars_20000.mat.pb -t sars_2000.nwk”.* We then generated pairwise alignment of each sequence with respect to *Wuhan-Hu-1 (NC_045512.2)* sequence (*ref.fa*, used as the reference sequence in UShER-MAT) using MAFFT, using the command: “*mafft --thread 32 --auto --keeplength --addfragments sars_20000.fa ref.fa > sars_20000.aln*”. We then used the generated alignment to produce variations for each sequence relative to the reference sequence (*Wuhan-Hu-1, NC_045512.2*) in a variant call format (VCF) file using *faToVcf* utility (http://hgdownload.soe.ucsc.edu/admin/exe/), using the command: “*faToVcf sars_20000.aln sars_20000.vcf*”. Finally, we reconstructed UShER-MAT using UShER program^30^, using the command: “*usher --threads 32 -t sars_20000.nwk --vcf sars_20000.vcf --save-mutation-annotated-tree sars_20000_.mat.pb*”. Once the UShER-MAT is reconstructed, the substitutions corresponding to each clade in the tree were extracted in a file (*clade.txt*) from UShER-MAT (*sars_20000.mat.pb*) using *matUtils*, using the command: “*matUtils extract -i sars_20000.mat.pb -C clades.txt*”.

To produce a PanMAN, we first generated multiple sequence alignment (MSA) of 20,000 SARS-CoV-2 sequences (*sars_20000.fa*, FASTA format) using MAFFT^62^ v7.520, using the command: “*mafft --thread 32 --nomemsave sars_20000.fa > sars_20000.msa*”, with default settings. Next we used *panmanUtils* to generate a PanMAT, single tree PanMAN, (*sars_20000.panman*) using the command: “*panmanUtils --msa-in sars_20000.msa --newick-in sars_20000.nwk*”, with the same tree topology as UShER-MAT. We extracted stored substitutions as well as indels corresponding to each clade in this PanMAT using the command: *“./panmanUtils -I sars_20000.panman --printMutations*”. To infer recombination, we used the RIVET pipeline^67^ to infer recombination events in the UShER-MAT (*sars_20000.mat.pb*). These inferred events, stored in a file (*cm.txt*), were then used to split the previously generated single-tree PanMAN into multiple trees interconnected with network edges in the final PanMAN (Fig. 4c) with *panmanUtils*, using the command: “.*/panmanUtils -I sars_20000.panman --complex-mutations cm.txt*”.

### System Details

All the experiments mentioned in this manuscript are performed on a 64-core Intel Xeon Silver 4216 processor, running at 2.1 GHz with 384 GB DDR4 RAM clocked at 3.2 GHz.

## Acknowledgments

We thank the authors and developers of the PanGraph tool, especially Marco Molari, and Liam P. Shaw, for their prompt and helpful responses to our inquiries and issues. We also thank Angie Hinrichs for her technical support and valuable feedback. Additionally, we acknowledge Cheng Ye and Carol Bao for technical support. We gratefully acknowledge all data contributors, i.e., the authors and their originating laboratories responsible for obtaining the specimens, and their submitting laboratories for generating the genetic sequence and metadata and sharing via the GISAID Initiative, on which this research is based. We are also thankful to the authors and laboratories that submitted genome data to NCBI and LANL databases, which helped this study. This work was supported by funding from the U.S. Centers for Disease Control and Prevention through the Office of Advanced Molecular Detection (CDC contract #75D30123C17463) and Amazon Research Award (Fall 2022 CFP).

## Data and Code Availability

All code developed in this study is freely available under the MIT license at https://github.com/TurakhiaLab/panman and the documentation is available at https://turakhia.ucsd.edu/panman. The PanMAN files produced during this study have been uploaded to an online repository available at https://zenodo.org/records/12630607.

## Supplementary Figures

**Supplementary Figure 1:**
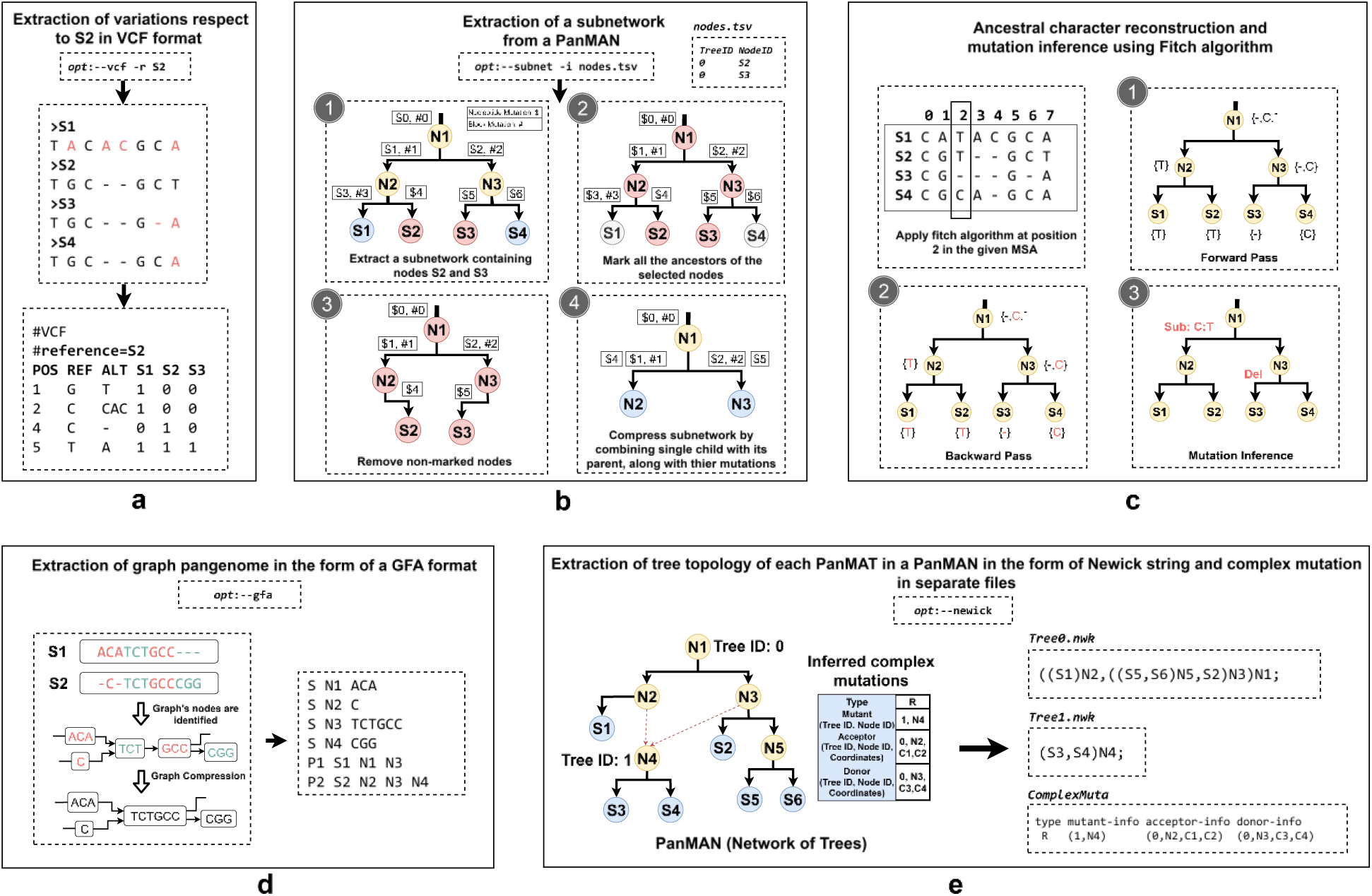
Functionalities in *panmanUtils*. Illustration of how *panmanUtils* a) extract variations with respect to any (S2 in this case) sequence from a PanMAT in a PanMAN, b) extract subnetworks from a network of trees in a PanMAN based on the provided list of node identifiers and corresponding tree identifiers in a TSV file, c) infers ancestral characters using Fitch algorithm, d) converts any PanMAT in a PanMAN into GFA file format, and e) extracts tree topology of every PanMAT in a PanMAN in the form of Newick string and complex mutation in separate files.

**Supplementary Figure 2:**
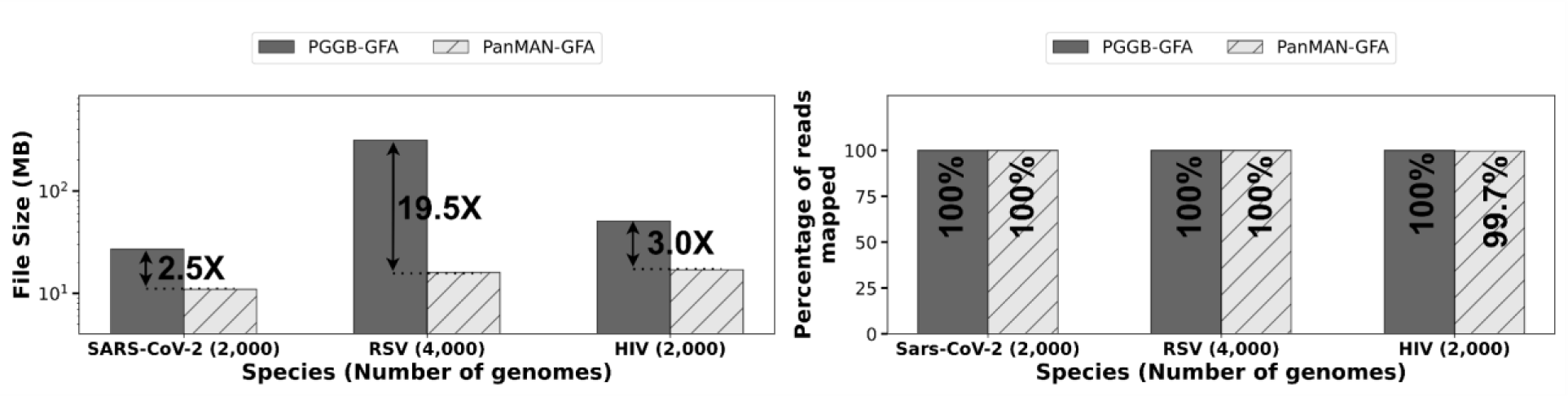
Evaluation of GFAs produced from PanMAN and compared against GFAs produced from PGGB pipeline. (a) Comparison of GFA file sizes produced from PanMAN and PGGB. (b) Comparison of percentage of reads mapped to GFAs produced from PanMAN and PGGB.

**Supplementary Figure 3:**
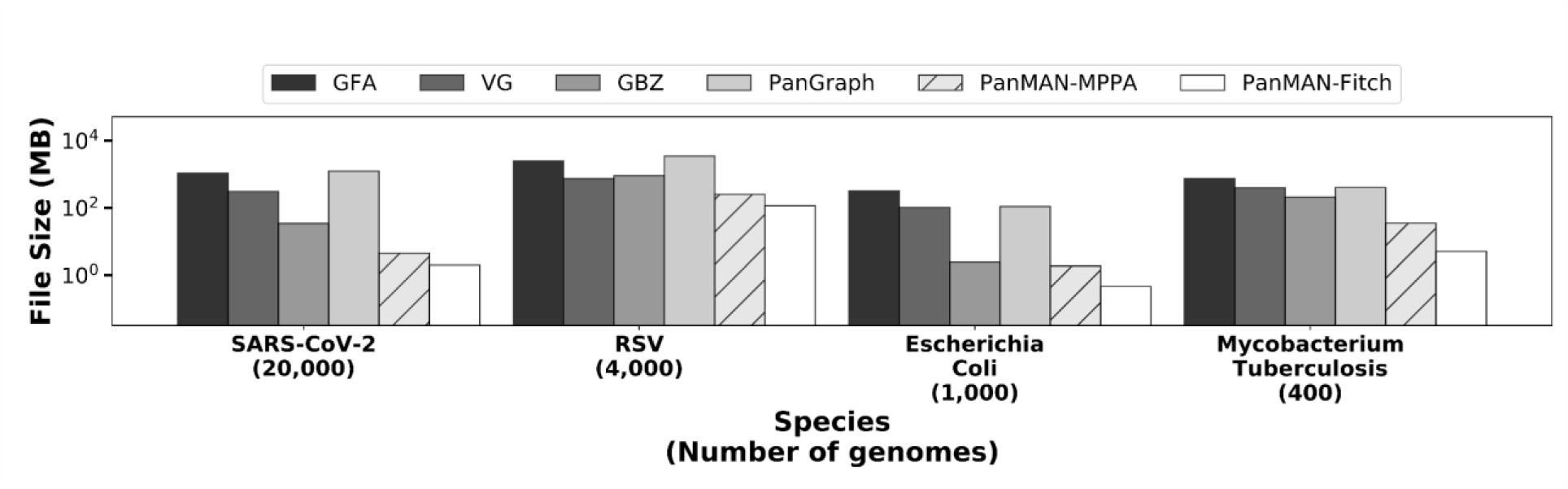
File sizes of PanMAN with different ACR methods -. Fitch (Parsimony-based) and MPPA (Likelihood-based), compared to other lossless alignment-preserving pangenomic formats.

**Supplementary Figure 4:**
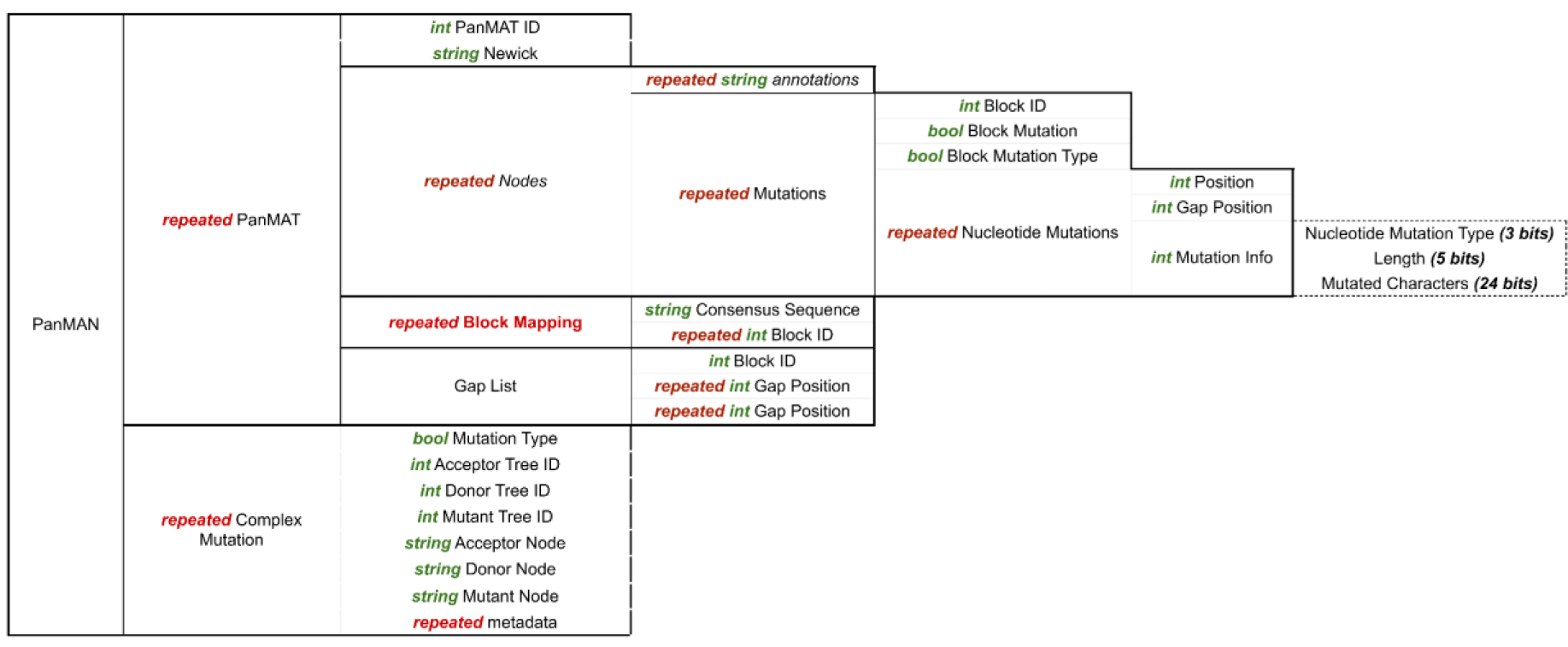
PanMAN protobuf file format: Hierarchy of the data structure that PanMAN utilizes for storing genomes using oogle’s protocol buffer (protobuf) serialization library.

**Supplementary Figure 5:**
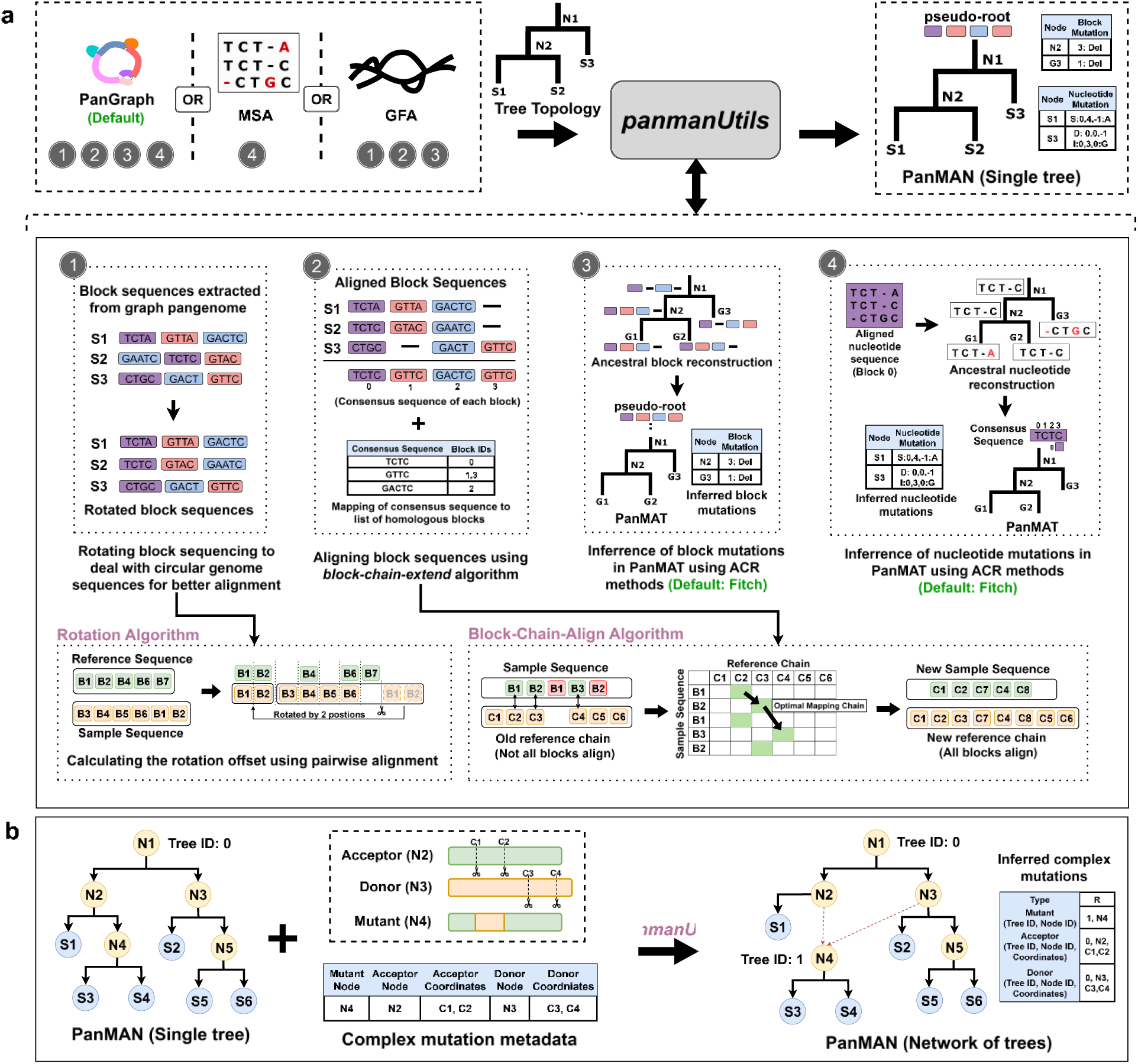
Overview of *panmanUtils* pipeline to construct PanMANs. PanMAN can be composed of a single or multiple PanMATs, a) illustrates *panmanUtils*’ input, output, and a detailed view of various steps involved in constructing a single tree PanMAN (or PanMAT), and b) illustrates how *panmanUtils*’ break a single tree into a network of multiple PanMATs using the inferred complex mutation metadata provided as input.

## Notes

### Competing Interest Statement

The authors have declared no competing interest.

### Summary of Updates

Figure 1 and Introduction have been revised

## References

1. Golicz, A. A., Bayer, P. E., Bhalla, P. L., Batley, J. & Edwards, D. Pangenomics Comes of Age: From Bacteria to Plant and Animal Applications. Trends Genet. 36, 132–145 (2020).

2. The Computational Pan-Genomics Consortium. Computational pan-genomics: status, promises and challenges. Brief. Bioinform. 19, 118–135 (2018).

3. Aggarwal, S. K. et al. Pangenomics in Microbial and Crop Research: Progress, Applications, and Perspectives. Genes 13, 598 (2022).

4. Shu, Y. & McCauley, J. GISAID: Global initiative on sharing all influenza data – from vision to reality. Eurosurveillance 22, (2017).

5. Clark, K., Karsch-Mizrachi, I., Lipman, D. J., Ostell, J. & Sayers, E. W. GenBank. Nucleic Acids Res. 44, D67–D72 (2016).

6. Rambaut, A. et al. A dynamic nomenclature proposal for SARS-CoV-2 lineages to assist genomic epidemiology. Nat. Microbiol. 5, 1403–1407 (2020).

7. De Bernardi Schneider, A., et al. SARS-CoV-2 lineage assignments using phylogenetic placement/UShER are superior to pangoLEARN machine-learning method. Virus Evol. 10, vead085 (2024).

8. Chen, C. et al. CoV-Spectrum: analysis of globally shared SARS-CoV-2 data to identify and characterize new variants. Bioinformatics 38, 1735–1737 (2022).

9. Obermeyer, F. et al. Analysis of 6.4 million SARS-CoV-2 genomes identifies mutations associated with fitness. Science 376, 1327–1332 (2022).

10. Tsui, J. L.-H. et al. Genomic assessment of invasion dynamics of SARS-CoV-2 Omicron BA.1. Science 381, 336–343 (2023).

11. Lam-Hine, T. et al. Outbreak Associated with SARS-CoV-2 B.1.617.2 (Delta) Variant in an Elementary School — Marin County, California, May–June 2021. MMWR Morb. Mortal. Wkly. Rep. 70, 1214–1219 (2021).

12. Li, T. et al. Genomic variation, origin tracing, and vaccine development of SARS-CoV-2: A systematic review. The Innovation 2, 100116 (2021).

13. Chalkias, S. et al. A Bivalent Omicron-Containing Booster Vaccine against Covid-19. N. Engl. J. Med. 387, 1279–1291 (2022).

14. Need, A. C. & Goldstein, D. B. Next generation disparities in human genomics: concerns and remedies. Trends Genet. 25, 489–494 (2009).

15. Brandt, D. Y. C. et al. Mapping Bias Overestimates Reference Allele Frequencies at the *HLA* Genes in the 1000 Genomes Project Phase I Data. G3 GenesGenomesGenetics 5, 931–941 (2015).

16. Günther, T. & Nettelblad, C. The presence and impact of reference bias on population genomic studies of prehistoric human populations. PLOS Genet. 15, e1008302 (2019).

17. Zhou, Y. et al. Graph pangenome captures missing heritability and empowers tomato breeding. Nature 606, 527–534 (2022).

18. Garrison, E. et al. Variation graph toolkit improves read mapping by representing genetic variation in the reference. Nat. Biotechnol. 36, 875–879 (2018).

19. Sherman, R. M. & Salzberg, S. L. Pan-genomics in the human genome era. Nat. Rev. Genet. 21, 243–254 (2020).

20. Heng Li. GFA-spec.

21. Sirén, J. & Paten, B. GBZ file format for pangenome graphs. Bioinformatics 38, 5012–5018 (2022).

22. Noll, N., Molari, M., Shaw, L. P. & Neher, R. A. PanGraph: scalable bacterial pan-genome graph construction. *Microb*. Genomics 9, (2023).

23. Li, H., Feng, X. & Chu, C. The design and construction of reference pangenome graphs with minigraph. Genome Biol. 21, 265 (2020).

24. Hickey, G. et al. Pangenome graph construction from genome alignments with Minigraph-Cactus. Nat. Biotechnol. (2023) doi:10.1038/s41587-023-01793-w.

25. Eggertsson, H. P. et al. Graphtyper enables population-scale genotyping using pangenome graphs. Nat. Genet. 49, 1654–1660 (2017).

26. Colquhoun, R. M., et al. Pandora: nucleotide-resolution bacterial pan-genomics with reference graphs. Genome Biol. 22, 267 (2021).

27. Rakocevic, G. et al. Fast and accurate genomic analyses using genome graphs. Nat. Genet. 51, 354–362 (2019).

28. Deorowicz, S., Danek, A. & Li, H. AGC: compact representation of assembled genomes with fast queries and updates. Bioinformatics 39, btad097 (2023).

29. Břinda, K. et al. Efficient and Robust Search of Microbial Genomes via Phylogenetic Compression. Preprint at 10.1101/2023.04.15.536996 (2023).

30. Turakhia, Y. et al. Ultrafast Sample placement on Existing tRees (UShER) enables real-time phylogenetics for the SARS-CoV-2 pandemic. Nat. Genet. 53, 809–816 (2021).

31. Kelleher, J., Thornton, K. R., Ashander, J. & Ralph, P. L. Efficient pedigree recording for fast population genetics simulation. PLOS Comput. Biol. 14, e1006581 (2018).

32. Richard R. Hudson. Gene genealogies and the coalescent process. (1990).

33. Kelleher, J. et al. Inferring whole-genome histories in large population datasets. Nat. Genet. 51, 1330–1338 (2019).

34. Speidel, L., Forest, M., Shi, S. & Myers, S. R. A method for genome-wide genealogy estimation for thousands of samples. Nat. Genet. 51, 1321–1329 (2019).

35. Schaefer, N. K., Shapiro, B. & Green, R. E. An ancestral recombination graph of human, Neanderthal, and Denisovan genomes. Sci. Adv. 7, eabc0776 (2021).

36. Zhang, B. C., Biddanda, A., Gunnarsson, Á. F., Cooper, F. & Palamara, P. F. Biobank-scale inference of ancestral recombination graphs enables genealogical analysis of complex traits. Nat. Genet. 55, 768–776 (2023).

37. glhaut, C., Pečerska, J., il, M. & Anisimova, M. Please Mind the ap: ndel-Aware Parsimony for Fast and Accurate Ancestral Sequence Reconstruction and Multiple Sequence Alignment including Long Indels. Preprint at 10.1101/2024.03.27.586611 (2024).

38. Ondov, B. D. et al. Mash: fast genome and metagenome distance estimation using MinHash. Genome Biol. 17, 132 (2016).

39. Li, H. et al. The Sequence Alignment/Map format and SAMtools. Bioinformatics 25, 2078– 2079 (2009).

40. Cochrane, G. et al. Facing growth in the European Nucleotide Archive. Nucleic Acids Res. 41, D30–D35 (2012).

41. Cardona, G., Rosselló, F. & Valiente, G. Extended Newick: it is time for a standard representation of phylogenetic networks. BMC Bioinformatics 9, 532 (2008).

42. Swofford, D. L. & Maddison, W. P. Reconstructing ancestral character states under Wagner parsimony. Math. Biosci. 87, 199–229 (1987).

43. Pupko, T., Pe, I., Shamir, R. & Graur, D. A Fast Algorithm for Joint Reconstruction of Ancestral Amino Acid Sequences. Mol. Biol. Evol. 17, 890–896 (2000).

44. Ree, R. H. & Smith, S. A. Maximum Likelihood Inference of Geographic Range Evolution by Dispersal, Local Extinction, and Cladogenesis. Syst. Biol. 57, 4–14 (2008).

45. Yang, Z. PAML 4: Phylogenetic Analysis by Maximum Likelihood. Mol. Biol. Evol. 24, 1586– 1591 (2007).

46. Oliva, A. et al. Accounting for ambiguity in ancestral sequence reconstruction. Bioinformatics 35, 4290–4297 (2019).

47. Ishikawa, S. A., Zhukova, A., Iwasaki, W. & Gascuel, O. A Fast Likelihood Method to Reconstruct and Visualize Ancestral Scenarios. Mol. Biol. Evol. 36, 2069–2085 (2019).

48. Huelsenbeck, J. P. & Bollback, J. P. Empirical and Hierarchical Bayesian Estimation of Ancestral States. Syst. Biol. 50, 351–366 (2001).

49. Pagel, M., Meade, A. & Barker, D. Bayesian Estimation of Ancestral Character States on Phylogenies. Syst. Biol. 53, 673–684 (2004).

50. Minh, B. Q. et al. IQ-TREE 2: New Models and Efficient Methods for Phylogenetic Inference in the Genomic Era. Mol. Biol. Evol. 37, 1530–1534 (2020).

51. Paul, P. et al. Genomic Surveillance for SARS-CoV-2 Variants Circulating in the United States, December 2020–May 2021. MMWR Morb. Mortal. Wkly. Rep. 70, 846–850 (2021).

52. Chen, Z. et al. Global landscape of SARS-CoV-2 genomic surveillance and data sharing. Nat. Genet. 54, 499–507 (2022).

53. Bloom, J. D., Beichman, A. C., Neher, R. A. & Harris, K. Evolution of the SARS-CoV-2 Mutational Spectrum. Mol. Biol. Evol. 40, msad085 (2023).

54. Ruis, C. et al. A lung-specific mutational signature enables inference of viral and bacterial respiratory niche. *Microb*. Genomics 9, (2023).

55. Boonsiri, T. et al. dentification and characterization of mutations responsible for the β-lactam resistance in oxacillin-susceptible mecA-positive Staphylococcus aureus. Sci. Rep. 10, 16907 (2020).

56. Karthikeyan, S. et al. Wastewater sequencing reveals early cryptic SARS-CoV-2 variant transmission. Nature 609, 101–108 (2022).

57. Li, X., Yan, H., Wong, G., Ouyang, W. & Cui, J. Identifying featured indels associated with SARS-CoV-2 fitness. Microbiol. Spectr. 11, e02269–23 (2023).

58. Hill, V. et al. The origins and molecular evolution of SARS-CoV-2 lineage B.1.1.7 in the UK. Virus Evol. 8, veac080 (2022).

59. Turakhia, Y. et al. Pandemic-scale phylogenomics reveals the SARS-CoV-2 recombination landscape. Nature 609, 994–997 (2022).

60. Zhan, S. H. et al. Towards Pandemic-Scale Ancestral Recombination Graphs of SARS-CoV-2. Preprint at 10.1101/2023.06.08.544212 (2023).

61. Roemer, C. et al. SARS-CoV-2 evolution in the Omicron era. Nat. Microbiol. 8, 1952–1959 (2023).

62. Katoh, K. MAFFT: a novel method for rapid multiple sequence alignment based on fast Fourier transform. Nucleic Acids Res. 30, 3059–3066 (2002).

63. Fitch, W. M. Toward Defining the Course of Evolution: Minimum Change for a Specific Tree Topology. Syst. Biol. 20, 406–416 (1971).

64. Huang, Y., Yang, C., Xu, X., Xu, W. & Liu, S. Structural and functional properties of SARS-CoV-2 spike protein: potential antivirus drug development for COVID-19. Acta Pharmacol. Sin. 41, 1141–1149 (2020).

65. Alisoltani, A., Jaroszewski, L., Iyer, M., Iranzadeh, A. & Godzik, A. Increased Frequency of Indels in Hypervariable Regions of SARS-CoV-2 Proteins—A Possible Signature of Adaptive Selection. Front. Genet. 13, 875406 (2022).

66. Resende, P. C. et al. The ongoing evolution of variants of concern and interest of SARS-CoV-2 in Brazil revealed by convergent indels in the amino (N)-terminal domain of the spike protein. Virus Evol. 7, veab069 (2021).

67. Smith, K., Ye, C. & Turakhia, Y. Tracking and curating putative SARS-CoV-2 recombinants with RIVET. Bioinformatics 39, btad538 (2023).

68. Goenka, S. D., Turakhia, Y., Paten, B. & Horowitz, M. SegAlign: A Scalable GPU-Based Whole Genome Aligner. in SC20: International Conference for High Performance Computing, Networking, Storage and Analysis 1–13 (IEEE, Atlanta, GA, USA, 2020). doi:10.1109/SC41405.2020.00043.

69. Gonnella, G., Niehus, N. & Kurtz, S. *GfaViz* : flexible and interactive visualization of GFA sequence graphs. Bioinformatics 35, 2853–2855 (2019).

70. Yuan, Y., Ma, R. K.-K. & Chan, T.-F. PanGraphViewer: A Versatile Tool to Visualize Pangenome Graphs. Preprint at 10.1101/2023.03.30.534931 (2023).

71. Miao, Z. & Yue, J.-X. VRPG: an interactive visualization framework for reference-projected pangenome graph. Preprint at 10.1101/2023.01.20.524991 (2023).

72. Wick, R. R., Schultz, M. B., Zobel, J. & Holt, K. E. Bandage: interactive visualization of *de novo* genome assemblies. Bioinformatics 31, 3350–3352 (2015).

73. Sanderson, T. Taxonium, a web-based tool for exploring large phylogenetic trees. eLife 11, e82392 (2022).

74. Kramer, A. M., Sanderson, T. & Corbett-Detig, R. Treenome Browser: co-visualization of enormous phylogenies and millions of genomes. Bioinformatics 39, btac772 (2023).

75. Katz, L. et al. Mashtree: a rapid comparison of whole genome sequence files. J. Open Source Softw. 4, 1762 (2019).

76. Yu, Y., Blair, C. & He, X. RASP 4: Ancestral State Reconstruction Tool for Multiple Genes and Characters. Mol. Biol. Evol. 37, 604–606 (2020).

77. Bouckaert, R. et al. BEAST 2: A Software Platform for Bayesian Evolutionary Analysis. PLoS Comput. Biol. 10, e1003537 (2014).

78. Armstrong, J. et al. Progressive Cactus is a multiple-genome aligner for the thousand-genome era. Nature 587, 246–251 (2020).

79. Garrison, E. et al. Building pangenome graphs. Preprint at 10.1101/2023.04.05.535718 (2023).

80. Maiolo, M. et al. ProPIP: a tool for progressive multiple sequence alignment with Poisson Indel Process. BMC Bioinformatics 22, 518 (2021).

81. Martin, D. P. et al. RDP3: a flexible and fast computer program for analyzing recombination. Bioinformatics 26, 2462–2463 (2010).

82. Li, X. et al. A Novel Strategy for Detecting Recent Horizontal Gene Transfer and Its Application to Rhizobium Strains. Front. Microbiol. 9, 973 (2018).

83. Adato, O., Ninyo, N., Gophna, U. & Snir, S. Detecting Horizontal Gene Transfer between Closely Related Taxa. PLOS Comput. Biol. 11, e1004408 (2015).

84. Michael, J. B., Robert, E. L. & Long Beach, C. Los Alamos Los Alamos National Laboratory Los Alamos, New Mexico 87545.

85. Pruitt, K. D., Katz, K. S., Sicotte, H. & Maglott, D. R. Introducing RefSeq and LocusLink: curated human genome resources at the NCBI. Trends Genet. 16, 44–47 (2000).

86. Voss, M., Asenjo, R. & Reinders, J. *Pro TBB: C++ Parallel Programming with Threading Building Blocks*. (Apress, Berkeley, CA, 2019). doi:10.1007/978-1-4842-4398-5.

87. Koranne, S. Boost C++ Libraries. in Handbook of Open Source Tools 127–143 (Springer US, Boston, MA, 2011). doi:10.1007/978-1-4419-7719-9_6.

88. Durbin, R., De Sanctis, B. & Blumer, M. Rotate: A command-line program to rotate circular DNA sequences to start at a given position or string. Wellcome Open Res. 8, 401 (2023).

89. Felsenstein, J. Evolutionary trees from DNA sequences: A maximum likelihood approach. J. Mol. Evol. 17, 368–376 (1981).

90. Byrd, R. H., Lu, P., Nocedal, J. & Zhu, C. A Limited Memory Algorithm for Bound Constrained Optimization. SIAM J. Sci. Comput. 16, 1190–1208 (1995).

91. Saitou, et al. The neighbor-joining method: a new method for reconstructing phylogenetic trees. Mol. Biol. Evol. (1987) doi:10.1093/oxfordjournals.molbev.a040454.

92. Li, H. Minimap2: pairwise alignment for nucleotide sequences. Bioinformatics 34, 3094– 3100 (2018).

93. Huang, W., Li, L., Myers, J. R. & Marth, G. T. ART: a next-generation sequencing read simulator. Bioinformatics 28, 593–594 (2012).

94. Sirén, J. et al. Pangenomics enables genotyping of known structural variants in 5202 diverse genomes. Science 374, abg8871 (2021).

95. McBroome, J. et al. A Daily-Updated Database and Tools for Comprehensive SARS-CoV-2 Mutation-Annotated Trees. Mol. Biol. Evol. 38, 5819–5824 (2021).

96. Turakhia, Y. et al. Stability of SARS-CoV-2 phylogenies. PLOS Genet. 16, e1009175 (2020).

